# Amyloid-beta and tau pathologies are both necessary to induce novel stage-specific microglia subtypes during Alzheimer’s disease progression

**DOI:** 10.1101/2021.10.16.464454

**Authors:** Dong Won Kim, Kevin Tu, Alice Wei, Ashley Lau, Anabel Gonzalez-Gil, Tianyu Cao, Kerstin Braunstein, Jonathan P. Ling, Juan C. Troncoso, Philip C. Wong, Seth Blackshaw, Ronald L. Schnaar, Tong Li

## Abstract

It is unknown whether specific microglia are selectively induced by amyloid-β (Aβ), tau pathologies, or both in combination. To address this, we use single-cell RNA-sequencing to profile mice bearing both Aβ and tau pathologies during Alzheimer’s disease (AD) progression. We identify novel microglia subtypes induced in a disease stage-specific manner. We show that during early-stage disease, interferon signaling induces a subtype of microglia termed EADAM. During late-stage disease, a second microglia subtype termed LADAM is detected. While EADAM and LADAM-like microglia are observed in other neurodegenerative models, the magnitude and composition of subtype markers are distinct from microglia observed with AD-like pathology. The pattern of EADAM- and LADAM-associated gene expression is observed in microglia from human AD, during the early and late stages of disease, respectively. Furthermore, we observe that several siglec genes are selectively expressed in either EADAM or LADAM. *Siglecg* is expressed in white-matter-associated LADAM, and expression of the human orthologue of *Siglecg* is progressively elevated in AD-stage-dependent manner but not shown in non-AD tauopathy. Our findings imply that both Aβ and tau pathologies are required for disease stage-specific induction of EADAM and LADAM.

## Introduction

Neuroinflammation is increasingly recognized as a key regulator of disease progression in neurodegenerative disorders^1,2^, including Alzheimer’s disease (AD), which is the most common cause of dementia^3^. Recent studies suggest that neuroinflammation serves as a mechanistic link between the development of amyloid-β (Aβ) plaques and tau neurofibrillary tangles, and the canonical pathologies of AD that are thought to drive synapse loss and neuronal death^3^. In addition to well-characterized AD susceptibility genes such as *APP, PSEN1 and PSEN2* (early-onset familial AD^4^), and *APOE* (late-onset AD^5,6^), several additional risk alleles for late-onset AD were identified in genes regulating immunomodulation, including *TREM2^7,8^*, phosphoinositide phospholipase C*γ*2 (*PLCG2*)^9^ and *CD33^10,11^.*

As the brain’s resident innate immune cells, microglia play multifunctional roles in brain health and the progression of neurodegenerative diseases such as AD^1,2,12^. Depending on the disease stage, microglia may protect against neurodegenerative proteinopathy and/or contribute to inflammatory damage^2,13^. Microglia maintain brain health by clearing cellular debris, including Aβ plaques and tau aggregates^13^. Microglial activation correlates positively with cognition and gray matter volume in humans, indicating that microglia can be protective, at least during early-stage AD^14^. However, microglia also secrete proinflammatory cytokines and can directly contribute to tau pathology^14,15^ and its subsequent neurotoxicity^1,2^. This dichotomous role of microglia in maintaining this balance between phagocytosis/clearance and pro-inflammatory mediator release is thought to be an essential and potentially targetable determinant of AD progression^16^.

Recent studies have identified numerous subtypes of microglia that display a dynamic range of responses and functions^17^, emphasizing the ability of microglia to serve a variety of important physiological roles^12^. Transcriptomic analyses of bulk tissues revealed disease-associated changes in microglia associated with AD^18,19^. Subsequent single-cell RNA-Sequencing (scRNA-Seq) approaches, however, were necessary to identify disease context-dependent microglial subtypes. scRNA-Seq has shed light on the spatial and developmental heterogeneity of microglia, and provides a high-resolution view of the transcriptional landscape of microglia subtypes during development and disease progression^17,20–23^. Since AD is a chronic disease with decades-long prodromal stages, understanding the disease stage-specific impacts of microglia subtypes is necessary to clarify the dual nature of microglia activation. The identification of disease-associated microglia (DAM), a unique TREM2-dependent subtype that expresses CD11c and is localized near Aβ plaques in an amyloidosis mouse model^20^, supports this notion. Despite these advances, a critical unresolved question is whether disease stage-specific microglia subtypes exist that are activated in response to both Aβ and tau pathologies. These microglia subtypes would represent novel therapeutic targets for modifying disease progression.

Here, we used an AD mouse model (*Tau4RΔK-AP* mice), in which wild-type tau is converted into tau aggregates to drive neuron loss in a neuritic plaque-dependent manner^24^. This animal model serves as an excellent mouse model to profile microglia subtypes across different stages of AD-like disease progression. To assess the requirement for both Aβ and tau pathologies to induce disease stage-specific microglia subtypes, control mice accumulating either Aβ plaques (*APP;PS1* mice) or tau tangles (*Tau4RΔK* mice) were also profiled. Using scRNA-Seq approaches, we found that microglia respond to the development of Aβ and tau pathologies in a disease stage-specific manner. During early-stage disease in 6-month-old *Tau4RΔK-AP*, but not *APP;PS1* or *Tau4RΔK* mice, the presence of both Aβ and tau pathologies triggered a novel microglia subtype we termed <u>E</u>arly-stage <u>AD</u>-<u>A</u>ssociated <u>M</u>icroglia (EADAM), which is distinct from DAM and express multiple interferon-regulated genes. We found that the signature genes in EADAM were also associated with a subgroup of microglia in the brains of early (Braak II). In late-stage disease (12-month-old *Tau4RΔK-AP* mice), we found that another novel microglia subtype emerged in response to tau pathology that we termed <u>L</u>ate-stage <u>AD</u>-Associated <u>M</u>icroglia (LADAM), and which expresses MHC and S100 family genes. We further observed a unique subtype of LADAM located near white matter, which is molecularly similar to a previously identified white matter-associated microglial subtype (WAM) that is increased during aging^25^, and undergoes additional molecular transitions with Aβ and tau pathologies.

LADAM microglia, including WAM-like LADAM, were observed in late (Braak VI), but not early (Braak II) stages of AD. Corroborating these findings, we found that *sialic acid-binding immunoglobulin-like lectin* (*Siglec*) family members are associated with specific subsets of microglia that are activated in a disease stage-specific manner in both mouse models of AD and in AD patients. For example, Siglec-F is upregulated in response to Aβ pathology and is selectively expressed in Aβ-associated DAMs, while Siglec-G is upregulated in white-matter-associated LADAM in late-stage AD. These findings are consistent with a model whereby both Aβ and tau pathologies are necessary to induce the emergence of EADAM and LADAM, and have important implications for the identification of novel molecular targets and therapeutic strategies for treatment of AD.

## Results

### ScRNA-Seq of microglia in mice harboring Aβ plaques and/or tau deposition

To identify microglia subtypes through AD-like pathology progression, we performed scRNA-Seq in the cerebral cortex and hippocampus of transgenic mice displaying Aβ plaques and/or tau pathologies. We took advantage of our previously characterized mouse model of AD (*Tau4RΔK-AP* mice), which exhibits AD-like pathologies including Aβ plaques and tau tangles, and result in progressive neuronal loss and brain atrophy^24^. In a cross-breeding strategy using mutant *APP;PS1* (AP)^26^ and *Tau4RΔK* mice, a cohort of *Tau4RΔK-AP* mice was generated. Female mice were aged to either 6-month-old or 12-month old. Six-month-old *Tau4RΔK* mice mimic an early AD stage, characterized by low levels of Aβ plaques (Fig.S1a,b) with minimum tau deposition in the hippocampus, and no loss of neurons. Twelvie-month-old *Tau4RΔK* mice mimic a late disease stage, characterized by robust Aβ plaques (Fig.S1c,d), tau deposition (Fig.S1e,f), and loss of neurons. For these two time points, in addition to cerebral cortices and hippocampi from *Tau4RΔK-AP* mice, we also collected littermate controls (*WT)*, *APP;PS1* (Aβ plaques), and *Tau4RΔK* (tau deposition and neuronal loss) mice and subjected these tissues across 4 genotypes to scRNA-Seq analysis.

We first analyzed female 6-month-old cerebral cortices and hippocampus, across 4 genotypes in triplicates (Fig.S2a). We observed many different cell types, including neurons, astrocytes, oligodendrocytes, and microglia (Fig.S2b,c). As previously shown^24^, we first confirmed the initiation of tau deposition in the absence of neuronal loss or brain atrophy in these mice (Fig.S2b, Fig.S4a), along with the presence of Aβ plaques (Fig.S1a,b). As expected, we observed a few DAM microglia in *Tau4RΔK-AP* brains in early-stage disease (Fig.S1g,h). Analyzing three biological replicate samples, we observed that microglia in *APP;PS1* and *Tau4RΔK-AP* mice (Fig.S1g,h, S2b,d, S4a), expressed a high level of disease-associated microglia 1 (DAM1) marker including *Cst7* (Fig.S2e). However, we failed to observe DAM2 markers, including *Gpnmb*, even in *APP;PS1* and *Tau4RΔK-AP* mice (Fig.S2f) at this age.

Compared to *Tau4RΔK* mice, Aβ plaques accelerated tau aggregation-dependent (Fig.S1c-f) neuronal loss and brain atrophy in 12-month-old *Tau4RΔK-AP* mice, as previously shown^24^. Using scRNA-Seq, we also identified cell clusters corresponding to major subtypes of neurons, glia, and immune cells (Fig.S3a-c) in 12-month-old mice of all four genotypes, analyzing three biological replicates. Consistent with our previous observation, AD-like pathologies in *Tau4RΔK-AP* mice led to gliosis and neuronal loss. An increase in the microglia population along with a corresponding decrease in the number of neurons was observed in the *Tau4RΔK-AP* mice (Fig.S1i,j, S3b,d, S4b). Compared to the other three control mice (*WT*, *APP;PS1,* and *Tau4RΔK* mice), *Tau4RΔK-AP* mice also showed an increase in immune cells, including genes that correspond to dendritic cells, natural killer cells, and neutrophils (Fig.1b). No dramatic changes in other defined non-brain-derived cell types were observed, with an exception of B cells and brain-associated macrophages in the *Tau4RΔK-AP* mice.

**Figure 1.**
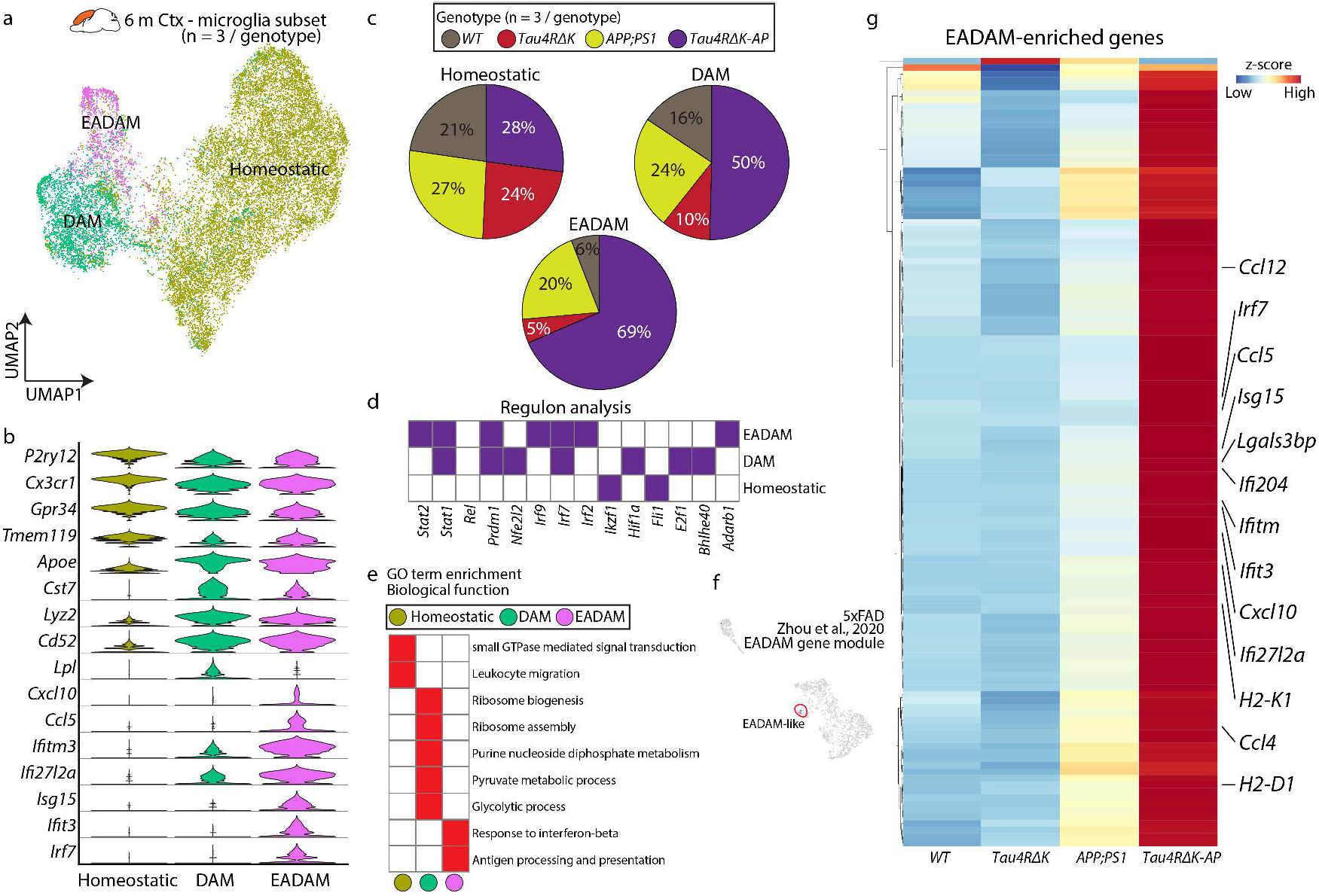
A unique microglia subtype EADAM is observed in the *Tau4RΔK-AP* mice in 6-month-old cortex. **a**, UMAP plot showing microglia clusters - Homeostatic, DAM, and EADAM, across 4 genotypes in 6-month-old cortex (n = 3/genotype). **b**, Violin plots showing top cluster genes of 3 microglia clusters. **c**, Pie graphs showing the distribution of 3 microglia clusters across genotypes (n = 3/genotype). **d**, Regulon analysis with SCENIC^62^, showing key regulons in each microglia cluster. **e**, GO analysis of top differential genes reveals a biological function in each microglial cluster. **f**, UMAP plot of sub-cluster of EADAM-like cluster in 7 month-old 5XFAD mice in Zhou et al 2020^23^. **g**, Heatmap showing expression of EADAM-enriched genes across genotypes.

Compared to *WT* and *Tau4RΔK* mice, both *APP;PS1* and *Tau4RΔK-AP* mice showed a marked increase in the proportion of microglia (Fig1i,j S4b). Known DAM1 marker genes such as *Cst7*, and DAM2 genes such as *Gpnmb*, were strongly expressed in both *APP;PS1* and *Tau4RΔK-AP* mice (Fig.S3e,f). These results establish that these scRNA-Seq datasets have sufficient quantitative power to identify microglia subtypes that are influenced by Aβ and/or tau deposition in a disease stage-specific manner.

### Identification of a novel <u>E</u>arly-stage <u>AD</u>-<u>A</u>ssociated <u>M</u>icroglia (EADAM) triggered by both Aβ plaques and tau deposition

We profiled microglia from the cerebral cortex and hippocampus of 6-month-old *Tau4RΔK-AP* mice, along with the three control mouse lines. Homeostatic microglia expressing genes such as *Tmem119* and *P2ry12*, were predominantly detected across all genotypes (Fig.1a), but two additional smaller microglia clusters were also observed (Fig.1a). The first microglia cluster expressed classic DAM1-like markers, including *Apoe* and *Cst7* (Fig.1b). The second microglia cluster expressed DAM1-like markers, but also displayed a unique gene module consisting of interferon-related genes, such as *Ifitm3, Ifit27l2a*, and *Ifit3* (Fig.1b). Since this cluster was predominantly composed of *Tau4RΔK-AP* microglia and detected during early-stage, presymptomatic stage of disease (Fig.1c), we thus identify this cluster as <u>E</u>arly-stage <u>AD</u>-<u>A</u>ssociated <u>M</u>icroglia (EADAM). Our regulon and pathway analysis confirmed the association of EADAM with interferon regulating transcription factor *Irf2/7/9* (Fig.1d) and linkage to the GO term ‘Response to Interferon-beta’ (Fig.1e).

More than 65% of EADAM were composed of *Tau4RΔK-AP* microglia, but approximately 20% of EADAM were composed of *APP;PS1* microglia (Fig.1c). EADAM-like microglia were also associated with a small sub-cluster in 7-month-old 5xFAD snRNA-Seq data^23^ (Fig.1f). We next compared differential gene expressions between EADAM populations. We identified that despite a significant contribution of *APP;PS1* microglia to EADAM cluster, the overall level of EADAM-enriched genes were expressed at a much lower level in *APP;PS1* microglia than that of *Tau4RΔK-AP* EADAM (Fig.1f). This observation is consistent with the previous report that accumulation of Aβ plaques leads to activation of the interferon pathway^27–30^. The expression of interferon pathway-related genes was robust in *Tau4RΔK-AP* microglia but was not observed in *Tau4RΔK* microglia, suggesting that both Aβ plaques and tau aggregation are necessary to fully induce EADAM differentiation. In summary, our data identify a novel microglia subtype termed EADAM that is triggered by a combination of Aβ plaques and tau deposition during early-stage AD.

### Identification of <u>L</u>ate-stage <u>AD</u>-<u>A</u>ssociated <u>M</u>icroglia (LADAM)

To further clarify the influence of Aβ plaques and tau tangles in microglia pathophysiology during late-stage AD, we unbiasedly subdivided microglia populations into five distinguishable clusters across all four genotypes (*WT*, *APP;PS1, Tau4RΔK*, and *Tau4RΔK-AP*) in 12-month-old mice. A few notable differences were observed in 12-month-old mice compared to 6-month-old mice. The DAM cluster was further divided into two clusters - DAM1 and DAM2. The DAM1 cluster expressed genes such as *Tyrobp, Ctsd, C1qa* (Fig.2b), whereas DAM2 expressed DAM1-enriched genes and additional disease-related genes, such as *Gpnmb*, *Cst7*, and *Spp1* (Fig.2b). The induction of DAM2 in response to Aβ plaques has been documented in a variety of APP-based mouse models of AD^20^. Indeed, both clusters were highly represented in both *APP;PS1,* and *Tau4RΔK-AP* mice (Fig.2c), but the DAM2 abundance in *Tau4RΔK* was low (Fig.2c). These observations are consistent with the view that these DAM clusters arise mainly as a result of Aβ plaques.

**Figure 2.**
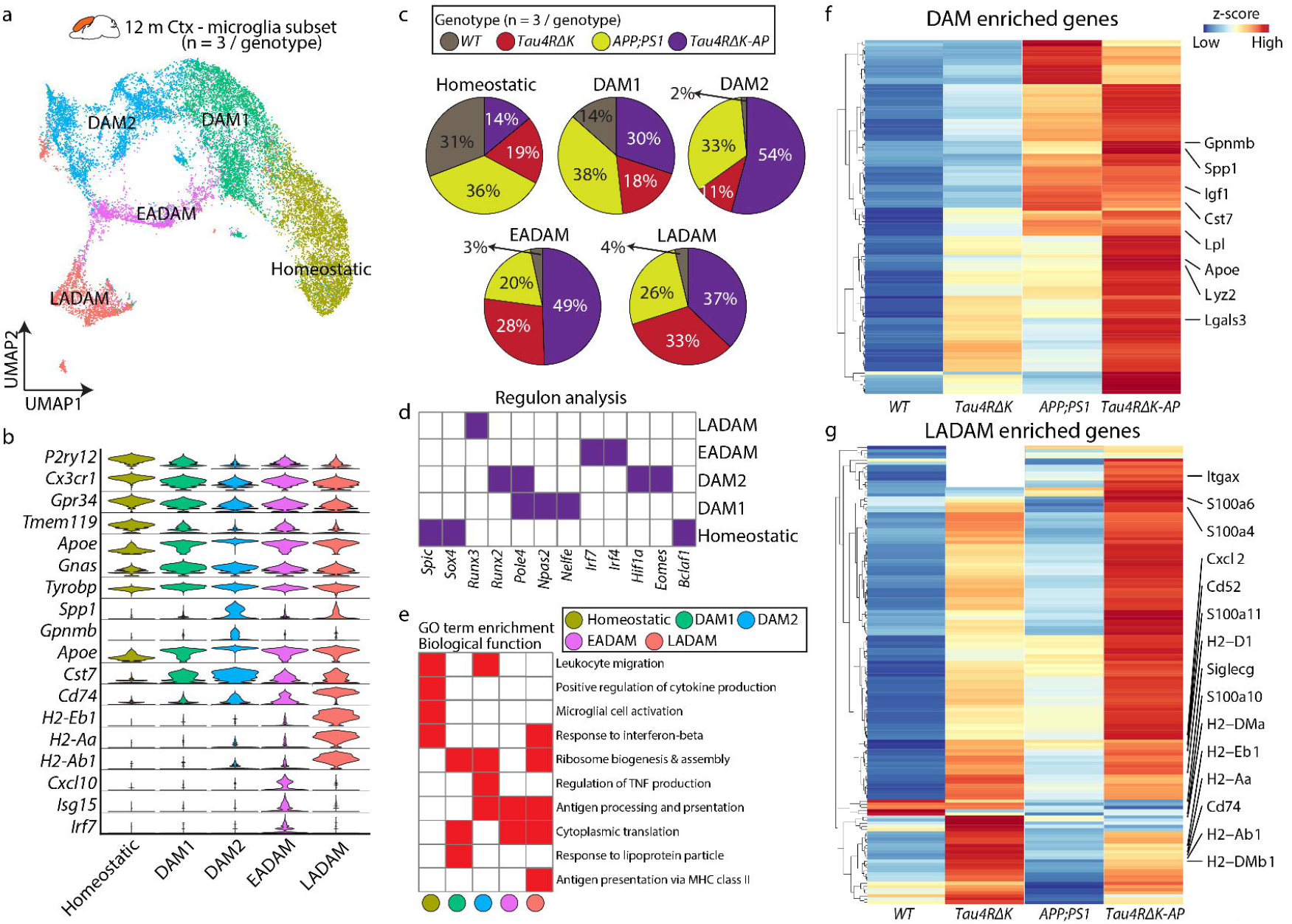
A unique microglia subtype LADAM is observed in the *Tau4RΔK-AP* mice in 12-month-old cortex. **a**, UMAP plot showing microglia clusters - Homeostatic, DAM1, DAM2, EADAM, and LADAM, across 4 genotypes in 12-month-old cortex (n = 3/genotype). **b**, Violin plots showing top cluster genes of 3 microglia clusters. **c**, Pie graphs showing the distribution of microglia clusters across genotypes (n = 3/genotype). **d**, Regulon analysis with SCENIC^62^, showing key regulons in each microglia cluster. **e**, GO analysis of top differential genes reveals a biological function in each microglia cluster. **f**, Heatmap showing expression of DAM-enriched genes across genotypes. **g**, Heatmap showing expression of LADAM-enriched genes across genotypes.

EADAM was still detected in 12-month-old *Tau4RΔK-AP* mice. Interestingly, EADAM-like microglia were also detected in *Tau4RΔK* mice, although they were absent at 6 month of age (Fig.2c). 12-month-old female *Tau4RΔK* mice already show some neuronal loss^24^, suggesting that while presence of tau aggregation alone may not trigger EADAM formation, cell death-induced inflammation may also trigger EADAM-like gene expression in microglia. We observed a unique microglia cluster that was not detected in 6-month-old mice (Fig.2a). Cells in this cluster expressed MHC Class II genes, such as *Cd74*, *H2-A2*, *H2-Eb1*, and *H2-Ab1* (Fig.2b). We termed these microglia <u>L</u>ate-stage <u>AD</u>-<u>A</u>ssociated <u>M</u>icroglia (LADAM). LADAM were found in *Tau4RΔK-AP* (37%), *APP;PS1* (26%), and *Tau4RΔK* (33%) samples (Fig.2c).

Regulon analysis identified key transcription factors associated with gene regulatory networks across all microglial clusters. Notably, while *Hif1a* and *Eomes* are selectively expressed in the DAM2, *Runx3* and *Irf4/7* are expressed in LADAM (Fig.2d), and GO analysis on LADAM shows ‘MHC Class II’ (Fig.2e). Similar to EADAM population in 6-month old mice, *Tau4RΔK-AP* mice generally have higher levels of DAM enriched genes (Fig.2f), indicating that while Aβ plaques can trigger DAM2, tau deposition and/or neuronal loss could further enhance DAM-enriched genes in *Tau4RΔK-AP* mice.

In addition, we found that many LADAM-specific genes (MHC Class II genes) are more highly expressed in *Tau4RΔK* than in APP*;PS1* microglia, suggesting that this microglia subtype might be induced by the development of tauopathies, such as tau tangles or tau phosphorylation. Moreover, relative to *Tau4RΔK* microglia, we found that LADAM gene signatures were more pronounced in *Tau4RΔK-AP* microglia (Fig.2g), which exhibited more robust tau pathologies. In addition to MHC class II genes, we also identified enriched expressions of *S100a* family genes, including *S100a4, S100a6*, and *S100a10* (Fig.2g).

These datasets show that tau pathologies are sufficient to trigger the emergence of LADAM and that their amplification is Aβ plaque-dependent. However, since MHC class II genes that are expressed in LADAM, such as *Cd74*, *H2-A2*, *H2-Eb1*, and *H2-Ab1* have also been previously observed in microglia in the CK-p25 neurodegeneration mouse model^31,32^ and also in some mouse models of late stage amyloidosis^27–29^, LADAM may be induced at least in part by neuronal death.

### Requirement of both Aβ plaques and tau deposition to elicit the emergence of disease stage-dependent microglia subtypes

To clarify how the emergence of EADAM, DAM2 and LADAM is regulated by Aβ plaques and tau deposition, we merged datasets obtained from microglia from both 6-month-old and 12-month-old samples (Fig.3a). RNA velocity analysis highlight that DAM1 is likely to give rise to, or to be closely related to, both DAM2 (Aβ plaques driven) and EADAM (Aβ plaque and tau deposition driven), whereas the convergence of DAM2 (Aβ plaque-driven and tau deposition-enhanced) and EADAM may ultimately lead to induction of LADAM (tau deposition/cell loss-driven and Aβ plaque-enhanced) (Fig.2b). As shown above, EADAM is composed mostly of cells derived from 6-month-old *Tau4RΔK-AP* mice (Fig.2b). The DAM2 and LADAM clusters were enriched in 12-month-old *APP;PS1*, *Tau4RΔK*, and *Tau4RΔK-AP* mice (Fig.2b).

**Figure 3.**
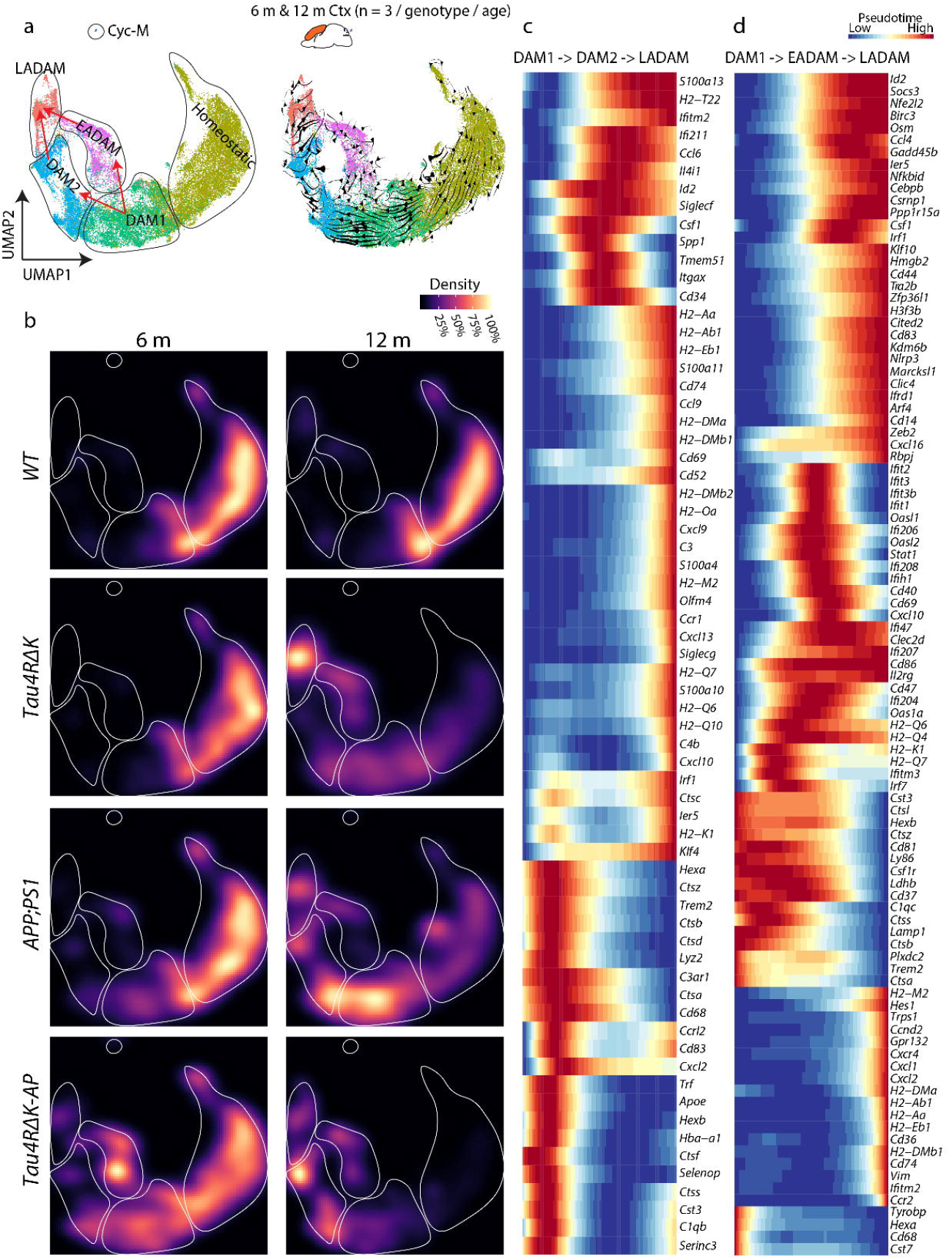
The emergence of disease stage-dependent microglia subtypes coincides with Aβ and tau deposition. **a**, UMAP plot showing microglia clusters in the 6- and 12-month-old cortex (all genotypes) (left) and UMAP plot with RNA velocity (right) showing 2 potential transitions between DAM1 and LADAM, with transitions into EADAM and/or DAM2 in between. **b**, UMAP plot showing the density of the captured microglia clusters across genotypes in the 12-month-old cortex. **c**, Pseudotime analysis from DAM1, DAM2, and LADAM. **d**, Pseudotime analysis from DAM1, EADAM and LADAM.

Furthermore, RNA velocity coupled with pseudotime analysis (Fig.4c,d) identified gene expression changes occurring during the temporal progression from DAM1 to DAM2 to LADAM, with changes in multiple inflammation-related genes, such as *Ccl6, Csf1, Cd34* (Fig.4c); as well as during the temporal progression from DAM1 to EADAM to LADAM, with changes in interferon genes, such as *Ifiti* gene families (Fig.4d). These data further support the previous findings that both Aβ plaques and tau deposition are critical for the induction of disease-stage specific microglia subtypes.

**Figure 4.**
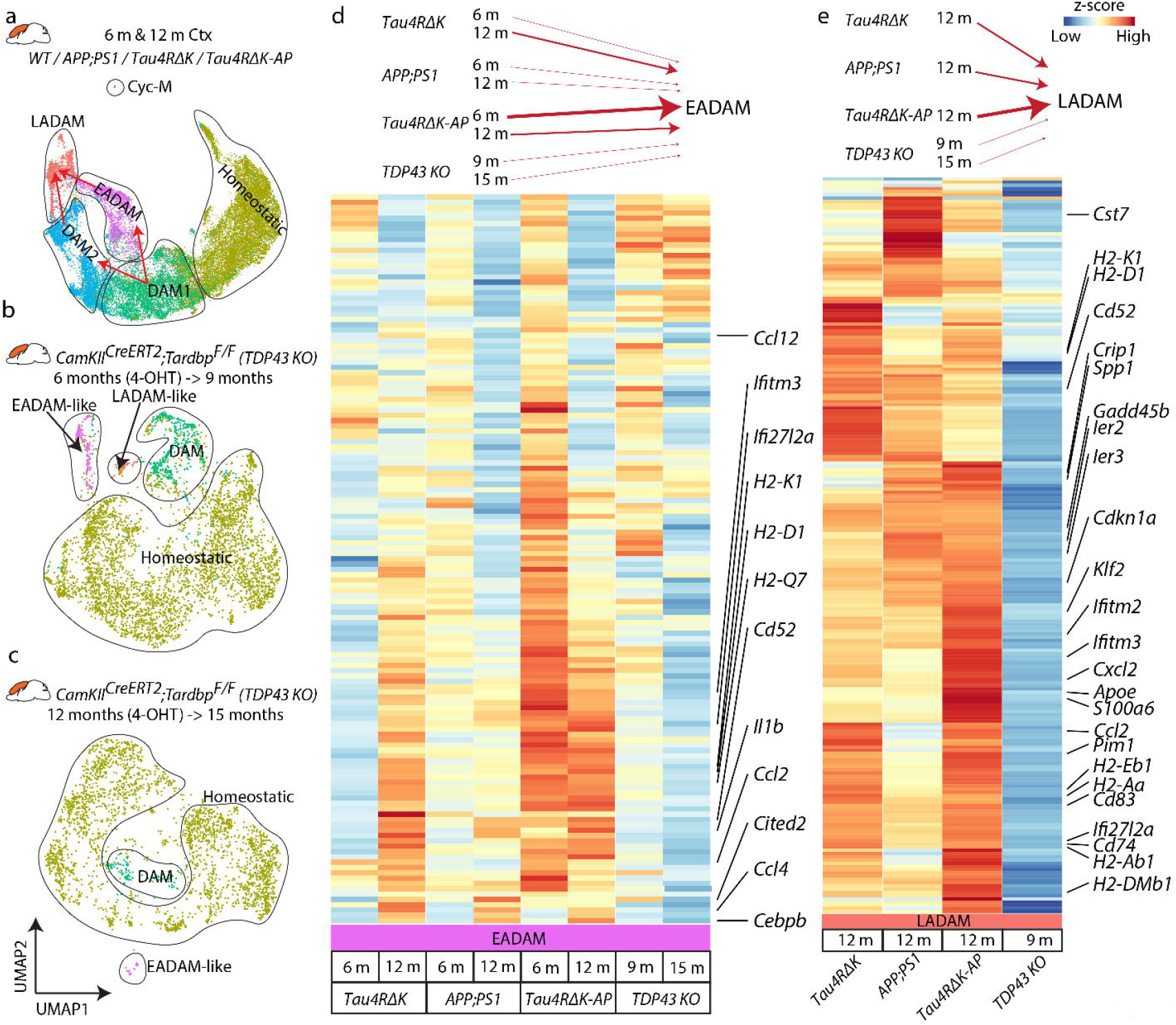
Analysis of EADAM and LADAM markers in other mouse models of neurodegenerative disease. **a**, UMAP plot showing microglia clusters in the 6 and 12-month-old cortex (all genotypes). **b**, UMAP plot showing microglia clusters in *CamKII^CreERT2^;Tardbp^lox/lox^* (TDP43 KO), 4-OHT inductio at 6 months and collected at 9 months. **c**, UMAP plot showing microglia clusters in *CamKII^CreERT2^;Tardbp^lox/lox^* (TDP43 KO), 4-OHT induction at 12 months and collected at 15 months. **d**, Heatmap plot showing expression of EADAM-enriched genes across AD genotypes and in TDP43 KO. **e**, Heatmap plot showing expression of LADAM-enriched genes across AD genotypes and in TDP43 KO.

A unique microglia cluster(s) that express either interferon genes and/or MHC Class II genes have been identified in other neurodegenerative models, such as CK-p25^32^ and in Trem2 deficient^33,34^ FTDP-17-linked tau model (P301L)^35^ crossed with PS2APP^36^ mice. This led to a hypothesis that potential conservation of EADAM- or LADAM-like subtypes might be shown across neurodegeneration and neuroinflammation.

To test this hypothesis, we then used another neurodegenerative mouse model *CamKII^CreERT2^;Tardbp^lox/lox^ (*TDP43 KO), where a tamoxifen induction lead to conditional deletion of TDP-43 in forebrain of the mice, resulting in selective vulnerability of hippocampal CA3 neurons^37^. We have generated scRNA-Seq data from the cerebral cortex and hippocampus of 2 different TDP43 KO samples, one being induced with 4-OHT at 6 months and collected at 9 months (TDP43 KO^6->9^) and the other being induced with 4-OHT at 12 months and collected at 15 months (TDP43 KO^12->15^) (Fig.4a-c). While both TDP43 KO mice showed DAM1-like clusters, TDP43 KO^6->9^ showed both EADAM- and LADAM-like clusters that respectively selectively express interferon genes or MHC Class II genes. However, TDP43 KO^12->15^ showed only EADAM-like clusters, with LADAM-like clusters not detected (Fig.4a-c).

We then integrated datasets from AD mice with those from TDP43 KO mice (Fig.4d,e). As mentioned above, both 6-month-old *APP;PS1* and *Tau4RΔK-AP* microglia showed a similar expression pattern of EADAM-enriched genes (Fig.4d), but *Tau4RΔK-AP* microglia displayed much higher expression levels of these genes than did *APP;PS1* microglia (Fig.4d). 12-month-old *Tau4RΔK* and *Tau4RΔK-AP* microglia showed overall lower expression levels, but broadly similar EADAM expression patterns (Fig.4d). Both TDP43 KO microglia showed a much weaker level of EADAM-enriched genes, similar to 6-month-old *APP;PS1* microglia, but TDP43 KO^6->9^ microglia much more closely resembled 6-month-old *APP;PS1* than TDP43 KO^12->15^ mice (Fig.4d). LADAM-enriched genes are composed of DAM2-enriched genes, MHC Class II genes, and *S100a* family genes. While, LADAM-like cluster in TDP43 KO^6->9^ microglia expressed MHC Class II genes, but the level of expression was much lower than any of *APP;PS1, Tau4RΔK* and *Tau4RΔK-AP* microglia (Fig.4e). We also failed to detect any DAM2 like clusters, or either *S100a* family genes and *Siglecg* in TDP43 KO^6->9^ microglia.

A similar observation was also made in CK-p25 microglia^32^(Fig.S5a). While integration of this dataset was not possible due to the use of different single-cell formats (SMART-Seq was used to generate the CK-p25 dataset), we noted that both EADAM-like interferon genes and LADAM-like MHC Class II genes were expressed higher in CK-p25 microglia than the control (Fig.S5b). However, both EADAM-like and LADAM-like microglia were intermingled, rather than forming separate clusters. We also failed to detect any *S100a* family genes and *Siglecg* expression in the dataset, as in TDP43 KO microglia (Fig.S5b).

This indicates that Aβ plaque or tau pathology independently induces specific gene signatures in microglia like EADAM and LADAM, and similar microglia clusters can also be observed in other neurodegeneration models. Neuroinflammation and/or cell loss with neurodegeneration diseases could be a potential trigger for microglia to develop EADAM- or LADAM-like gene expression profiles. However, the combination of both Aβ plaque and tau pathology result in more EADAM and LADAM, much higher expression levels of the signature genes, and induction of additional molecular markers in LADAM.

### Identification of stage-specific EADAM and LADAM signatures in Alzheimer’s Disease

We next generated snRNA-Seq dataset from the human entorhinal cortex (ERC) at Braak stage 2, where AD pathology is first detected and shares a pathological resemblance to our 6-month-old *Tau4RΔK-AP* mice (Fig.5a-c). We also generated snRNA-Seq from the superior frontal gyrus (SFG) at Braak stage 2, 4 and 6 (Fig.5a-c). Braak stage 2 SFG is devoid of any AD pathology, and Braak stage 4 and 6 SFG represent late stages of AD, and shares pathological resemblance to the 12-month-old *Tau4RΔK-AP* mice.

**Figure 5.**
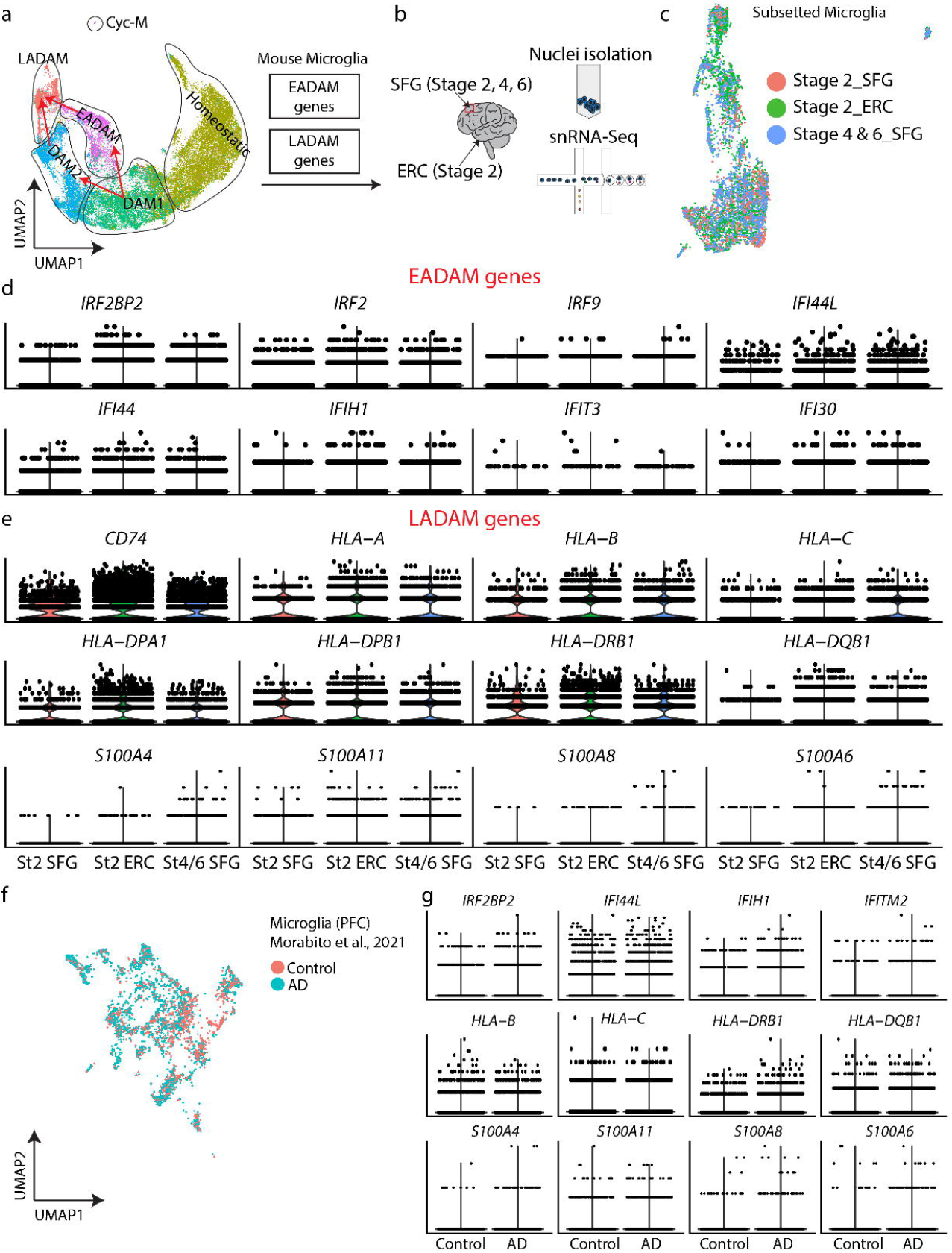
6-month-old EADAM at and 12-month-old LADAM clusters in *Tau4RΔK-AP* miceat resemble microglial subtypes seen in snRNA-Seq from AD samples. **a**, UMAP plot showing microglia clusters in the 6 and 12-month-old cortex (all genotypes) from Fig 4a. EADAM and LADAM-enriched gene homologs were looked at in human snRNA-Seq. **b**, Schematic showing snRNA-Seq from human postmortem samples, Stage 2 entorhinal cortex (ERC), superior frontal gyrus (SFG), Stage 4 SFG, and Stage 6 SFG. **c**, UMAP plot showing microglia from human snRNA-Seq. **d**, Violin plots showing expression of EADAM-like interferon genes. **e**, Violin plots showing expression of LADAM-like MHC and S100 gene families. **f**, UMAP plot showing microglia from human snRNA-Seq dataset from Morabito et al., 2021. **g**, Violin plots showing IFN, MHC and S100 genes in the dataset from Morabito et al., 2021.

We initially checked the expression pattern of EADAM- and LADAM-enriched gene homologs in human snRNA-Seq dataset (Fig.S6), and observed a moderate conservation in gene expression pattern. Most EADAM-enriched genes were seen at Braak stage 2 SFG and ERC (Fig.S6a), while LADAM-enriched genes were seen at Braak stage 4 and 6 SFG (Fig.S6b). Some inconsistency of expression patterns might be due to the absence of AD-specific microglia clusters in the human snRNA-Seq dataset, despite having a high resolution of 6,000 cells. This observation has been previously shown in other human snRNA-Seq dataset in AD or other neurodegeneration ^38–40^. The difference might be due to the fact that AD and other neurodegenerative disease typically progress slowly, over many years, and all microglia populations may end up transitioning to disease-associated states. This timing is different in virtually all animal models that were designed to undergo neurodegeneration much more rapidly.

While the presence of EADAM- and LADAM-like microglial subtypes in AD demonstrated important similarities with our mouse models, we also wanted to analyze expression of MHC and interferon-induced genes. We observed a consistent changes in interferon genes, such that transcriptional factor/co-regulator *IRF2BP2*, *IRF2*, and *IRF9* showed higher expression at Braak stage 2 ERC than SFG, and expression levels of these genes were decreased in Braak stage 4/6 SFG (Fig.5d). There also was a similar trend in expression of other interferon-regulated genes, including *IFI44*, *IFI44L* (Fig.5d). We also observed a similar trend in MHC genes, such as *HLA-A*, *HLA-B*, and *HLA-DPA1*, which showed a higher expression level in Braak stage 2 ERC and Braak stage 4/6 SFG than Braak stage 2 SFG (Fig.5e). More importantly, *S100* genes, such as *S100A4*, *S100A11*, were exclusively detected in Braak stage 4/6 SFG (Fig.5e). This observation was also conserved in other high-quality human AD snRNA-Seq from prefrontal cortex^38^(Fig.5f-g), where interferon-regulated, MHC, and *S100* family genes were more highly expressed in AD samples than in the control (Fig.5f-g).

These findings thus establish the emergence of EADAM-related genes in early AD-stage and LADAM-related genes in late AD-stage, and suggest that these microglia subtypes are induced by both Aβ plaques and tau deposition in a disease stage-specific manner.

### Siglecs serve as specific biomarkers to track Alzheimer’s disease stage-specific microglia subtypes

Our analysis identifies multiple microglial genes that are selectively expressed in late-stage AD. To further characterize late-stage AD-specific genes, we focused on the Siglec gene family, several of which are thought to be relevant for AD progression^41^. Homeostatic microglia expressed *Mag*, *Siglece*, *Cd33*, *Siglech* (Fig.6a,b), EADAM expressed similar Siglec genes as homeostatic microglia, but also expressed *Siglec1* (Fig.6a,b). Both DAM2 and LADAM clusters expressed high levels of *Siglecf* (Fig.6a,b). Both *Siglec1* and *Siglecg* were enriched in the LADAM. *Siglecf*, in particular, was enriched in both *APP;PS1* and *Tau4RΔK-AP* mice (Fig.6c, Fig.S6a) and showed increased expression in late-stage disease (Fig.6d, Fig.S6b).

**Figure 6.**
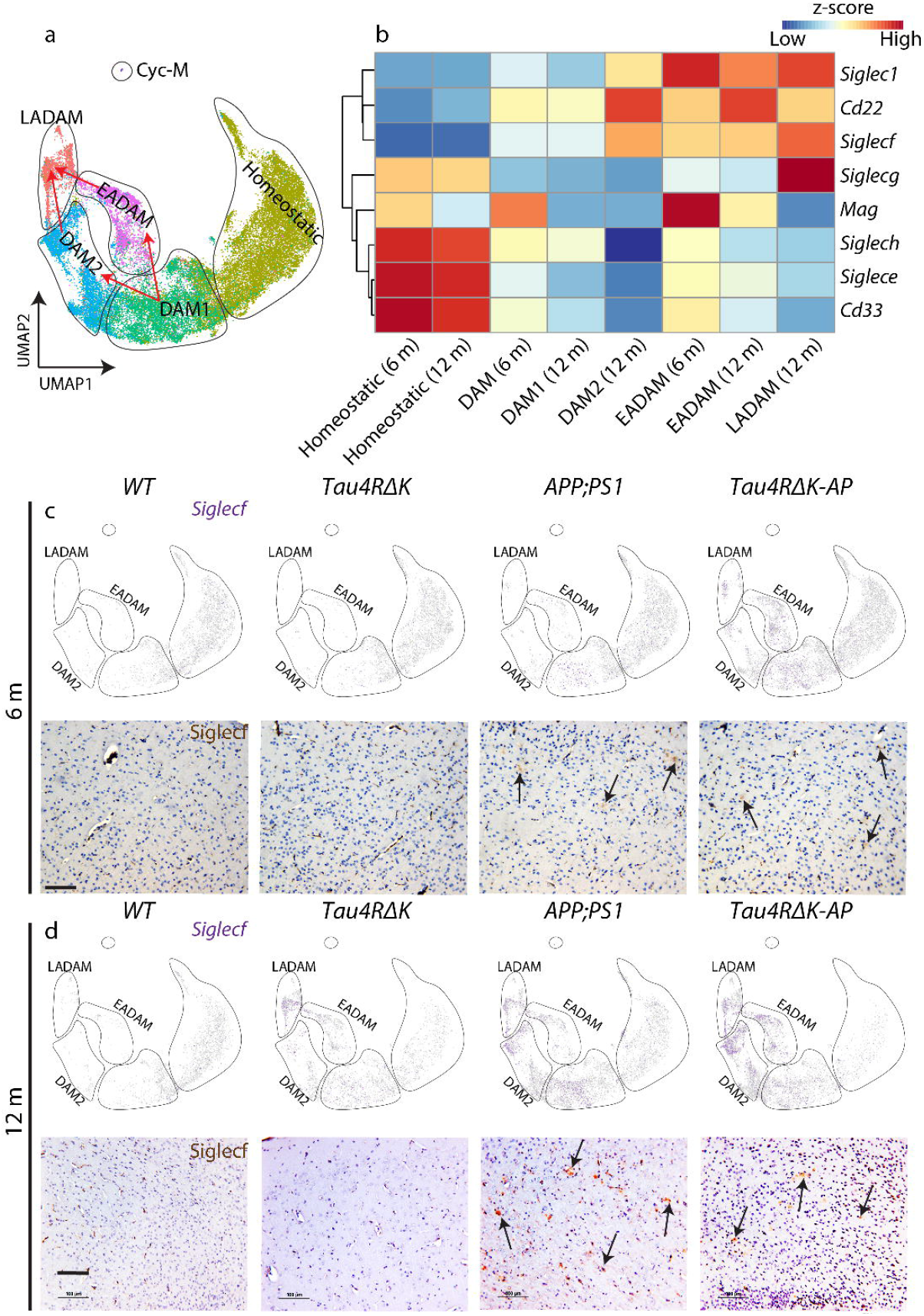
Siglec-genes show cluster- and genotype-specific expression patterns. **a**, UMAP plot showing microglial clusters in the 6 and 12-month-old cortex (all genotypes). **b**, Heatmap plot showing Siglec genes expressions across microglial clusters. **c, d**, UMAP plot showing *Siglecf* expression in *WT*, *Tau4RΔK*, *APP;PS1*, *Tau4RΔK-AP* at 6-month-old (c) and 12-month-old (d) (top). Note a higher expression of *Siglecf* in DAM2 in *APP;PS1* and *Tau4RΔK-AP* mice at 12-month-old. Siglec-F immunostaining was performed in the cortex in *WT*, *Tau4RΔK*, *APP;PS1*, *Tau4RΔK-AP* at 6-month-old (c) and 12-month-old (d) (bottom). Scale bars = 100 μm.

Some LADAM markers, particularly MHC II genes, were expressed in other neurodegenerative models (Fig.5, S6), but *S100* family of genes and *Siglecg* was unique to late-stage AD microglia in both animal models and in human late stage AD snRNA-Seq. This implies that induction of the LADAM subcluster is Aβ- and tau-dependent. We further identified 4 subclusters within the LADAM cluster (Fig.7a). Sub-clusters 2 and 3, in particular, were most robustly and selectively expressed in *Tau4RΔK-AP* mice (Fig.7b).

**Figure 7.**
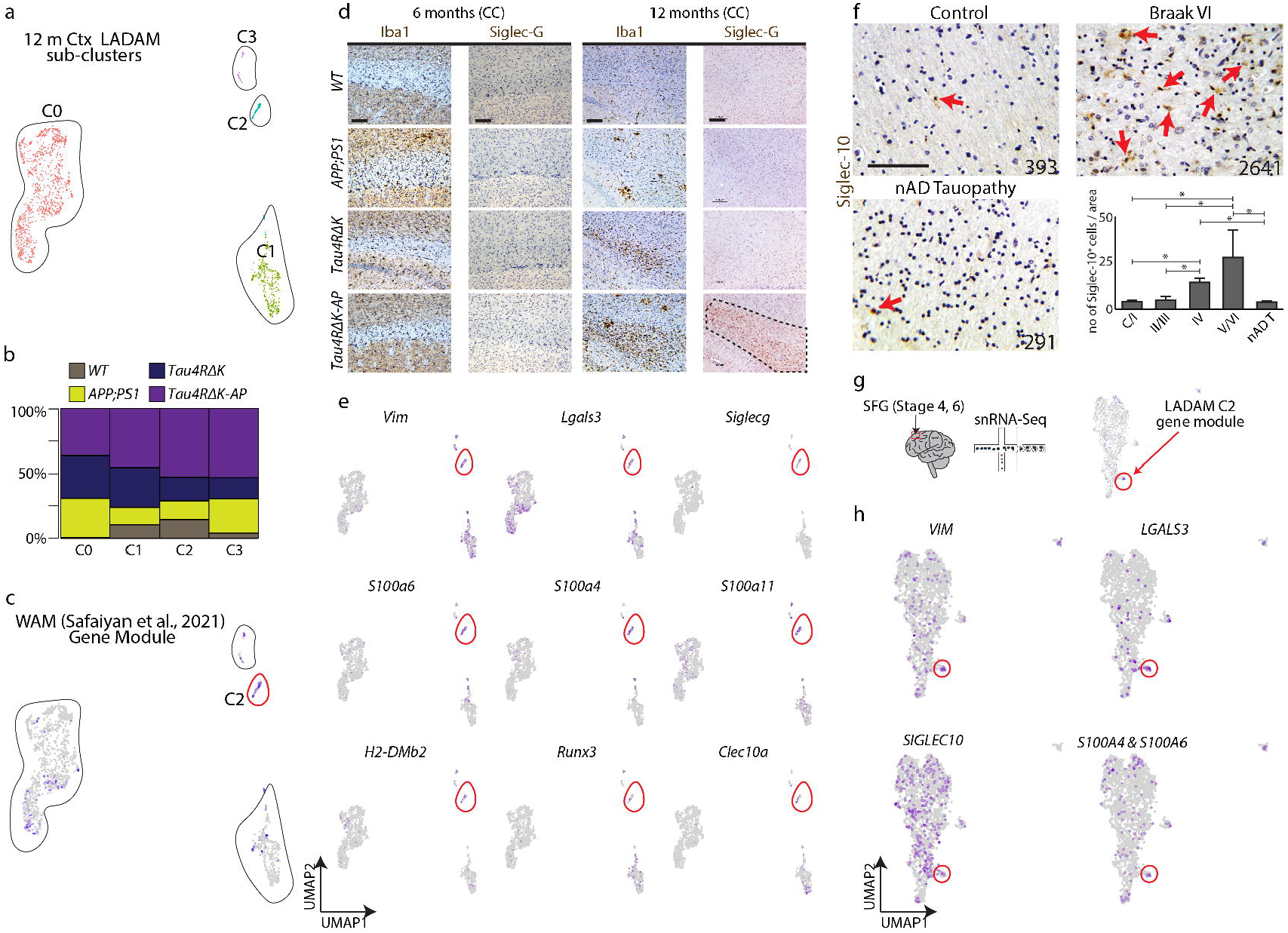
A LADAM subcluster is derived from white-matter-associated microglia. **a**, UMAP plot of 12-month-old ‘LADAM’. **b**, Bar plot showing the distribution of 4 genotypes across 4 LADAM clusters. **c**, UMAP plot of LADAM, showing cells/clusters (purple colored cells) that resemble WAM^25^. Note that C2 has the higher enrichment with WAM gene modules. **d**, Iba1 and Siglec-G immunostaining in 6-month-old and 12-month-old corpus callosum (CC) in *WT*, *APP;PS1*, *Tau4RΔK*, and *Tau4RΔK-AP*. **e**, UMAP plot showing gene expressions that are highly enriched in the *Tau4RΔK-AP* enriched (LADAM) WAM cluster. **f**, Siglec-10 immunostaining in human cortex samples in control, Braak stage 6, and nAD tauopathy (numbers on figure panels indicate BRC#, immunostaining of other Braak stages are shown in Fig.S7), with bar graphs showing quantification of Siglec-10 cells in white matter. **g**, Schematic showing human snRNA-Seq on Stage 4 and 6 SFG (left) and UMAP plot showing LADAM C2 enriched gene module. **h**, UMAP plot showing expression of LADAM C2 cluster markers in human snRNA-Seq. Scale bars = 100 μm.

Cluster 2, a LADAM sub-cluster that is marked by *Siglecg*, also exhibited genes previously shown to be expressed in <u>W</u>hite matter-<u>A</u>ssociated <u>M</u>icroglia (WAM), such as *Lgals3*, *Fam20c*, *Vim* (Fig.7a,c)^25^. In WAM-like LADAM, we observed that this cluster also expressed *S100a6*, *S100a4, S100a11, H2-DMb2, Runx2,* and *Clec10a* (Fig.7e). We also observed a small cluster that resembles WAM from the 6-month-old mouse cortex (Fig.S7f), but this small population did not express any LADAM markers in the 12-month-old cortex. We did not detect Siglec-G protein in the 6-month-old corpus callosum or hippocampus (Fig.7d). Siglec-G immunoreactivity was robust in the corpus callosum, and increased in *Tau4RΔK-AP* microglia (Fig.2d), as well as in the cortex at 12 months (Fig.S7e). However, we failed to observe Siglec-G immunoreactivity in *APP;PS1* or *Tau4RΔK* microglia at the same age, despite increased microglia populations near the corpus callosum (Fig.7d).

To corroborate this finding in AD, we used antisera directed against Siglec-10 (the human homolog of Siglec-G)^42^ to screen AD brains at multiple Braak stages. We found Braak stage-dependent accumulation of Siglec-10 in AD brains (Fig.7f, S8), with Siglec-10 signal was significantly increased from Braak stage 4 (Fig.7f, S8), but Siglec-10 signal was not increased in non-AD tauopathy (Fig.7f, S8). To further validate these observations, we checked snRNA-Seq on human postmortem Braak stage 4 and 6 SFG (Fig.7g). As mentioned above, late-stage AD microglia (Braak stage 4 and 6 SFG), expressed high levels of *CD74* and other LADAM genes (Fig.5). We also observed a small population of WAM-like LADAM, as seen in 12-month-old *Tau4RΔK-AP* mice (Fig.7g), such as *SIGLEC10, S100A6, LGALS3, FAM20C* (Fig.7g). Siglecs, in particular Siglec10, is disease-specific, as the level of Siglec-10+ cells in non-AD tauopathy was similar to that of control and low Braak stage AD (Fig.7f, S8), indicating that Siglec-10 serves both an AD-specific biomarker and a potential therapeutic target associated with late-stage AD.

## Discussion

### Discovery of novel disease stage-specific microglial subtypes induced by Aβ plaques and tau pathology

The identification of multiple AD risk alleles implicated in the regulation of inflammation^13,43^ and of DAM induction in response to Aβ plaques^20^, strongly support the hypothesis that microglia-driven neuroinflammation is a key regulator of AD progression. However, critical information regarding microglia subtypes that respond to disease stage-specific canonical AD pathologies of Aβ and tau, particularly during early disease stages, remains elusive. To address this, we took advantage of our *Tau4RΔK-AP* mouse model, which exhibits both Aβ plaques and tau deposition, and profiled microglia subtypes during disease progression using scRNA-Seq. To tease apart the influence of Aβ plaques from that of tau deposition, we also profiled microglial subtypes in *APP;PS1* (model of Aβ plaques) and *Tau4RΔK* (model of tau deposition) littermate mice. We identified several disease stage-specific subtypes of microglia that are selectively induced in response to Aβ and/or tau.

We discovered the emergence of another novel microglia subtype, <u>E</u>arly-stage <u>AD</u>-<u>A</u>ssociated <u>M</u>icroglia (EADAM), that is selectively induced by both Aβ and tau pathologies but not either Aβ or tau pathologies alone, during early-stage disease before AD-like pathologies are observed. EADAM are characterized by many interferon-pathway-related genes (*Ifit3, Ifit204, Ifit2*) and interferon regulating transcription factors (*Irf2/7/9)* as well as inflammatory chemokines (*Ccl12, Cxcl10, Ccl5, Ccl2*). Some of the genes in EADAM have been observed in other mouse models of amyloidosis that show early AD-like pathology^27–29^, including *APP;PS1* and 5xFAD. However, given that EADAM-enriched genes are expressed in much higher levels in our *Tau4RΔK-AP* microglia, this strongly supports the view that both Aβ and tau pathologies are required to induce disease stage-specific microglia subtypes in AD.

We also identified previously described DAM in both mouse lines in which Aβ plaques are induced (*APP;PS1* and *Tau4RΔK-AP*)^20,44^. However, DAM induction in *Tau4RΔK* mice was blunted, supporting the hypothesis that DAM are primarily induced by Aβ plaques and/or inflammation, but not by tau pathology. While Aβ plaques alone are sufficient to induce DAM2, this subtype can be greatly amplified by both Aβ and tau pathologies, giving rise to LADAM. LADAM continues to express classic DAM genes but also upregulates additional genes, including *Cd74* and MHC Class II genes. These broad LADAM marker-expressing microglia (*Cd74* and MHC Class II genes) have been previously shown in some amyloidosis mouse models at late stages (known as activated response microglia)^27–29^, but these genes are likely to be contributed by mild tau phosphorylation shown in these amyloidosis mouse models, based on our findings from *Tau4RΔK* (Fig.2). In addition, since LADAM-like clusters and markers were observed in other neurodegeneration models, such as TDP43 KO and CK-p25^31,32^, LADAM-enriched genes might be triggered by neurodegeneration and/or inflammation in addition to tau pathologies. However, much like EADAM, LADAM-enriched genes were expressed in *Tau4RΔK-AP* microglia, indicating that the full set of LADAM-specific genes is triggered by the combination of both Aβ and tau pathologies. Furthermore, the combination of Aβ and tau pathologies induced LADAM to express unique *S100a* genes and *Siglecg*.

In particular, a sub-cluster of LADAM that are localized near white matter tracts during late-stage disease shares a similar gene signature with previously identified <u>w</u>hite-matter-<u>a</u>ssociated <u>m</u>icroglia (WAM). WAM have been previously shown to be present in 24 month-old wildtype mice^25^. Indeed, microglia near white matter tracts were detected in all 4 genotypes in 12-month-old mice, but only mice with Aβ plaques and tau deposition showed expression of specialized LADAM gene signatures such as *Siglecg* and *S100a6.* Aβ plaques and tau deposition induce new molecular signatures in previously identified WAM, leading to the formation of this LADAM subcluster.

While our discovery of novel microglia subtypes has important clinical implications, the mechanisms by which EADAM and LADAM emerge during different stages of disease progression in response to Aβ plaques and tau deposition remain to be fully established, but our current study provides a potential regulators of these microglia populations. Neuronal loss induced by aging and/or tau pathologies may play an essential role in regulating this process.

### Identifications of disease stage-specific microglia subtypes in AD

Emergence of EADAM during presymptomatic AD stage and detection of LADAM when AD-like pathologies are visible led to the hypothesis that AD brains may harbor similar unique microglia clusters across disease progression. EADAM were observed in the entorhinal cortex during early (Braak II) but not late (Braak VI) stage disease, strongly supporting the view that EADAM first emerge in response to initial stages of Aβ plaque formation and tau deposition, corresponding to a stage of negligible or mild cognitive impairment. The microglia cluster expressing IFN-related gene that is similar to EADAM, has been shown in the FTDP-17-linked tau model (P301L)^35^ crossed with PS2APP^36^, but only when *Trem2* is deficient ^33,34^. The FTDP-17-linked tau model (P301L) can result in mild tau aggregation independent of Aβ plaques, but our model requires Aβ plaques to drive the pathological conversion of wild-type mouse tau to form tau tangle-like aggregates. Since *Trem2* deficiency can result in the spreading of tau aggregates^45^, the previous finding^34^ aligns well with our finding that Aβ plaques and tau deposition are required to give rise to EADAM.

In contrast, the emergence of LADAM in both the entorhinal cortex and superior frontal gyrus did not occur until Braak IV stage, implying that these subtypes are induced in response to more severe Aβ and tau pathologies accompanied by neuronal loss during late-stage disease. Some LADAM-enriched genes such as *HLA-DRB1* and *HLA-DRB5* positively correlate with AD pathology^46^, and share a few similar molecular signatures (e.g. MHC Class II genes) with lipid droplet-accumulating microglia^47^. We also observed an upregulation of *S100A* genes in late stage AD.

The fact that EADAM are first detected in 6-month-old *Tau4RΔK-AP* mice and Braak stage II AD datasets, while LADAM are first detected in 12-month-old *Tau4RΔK-AP* mice and Braak stage IV AD datasets, demonstrates that disease stage-dependent changes in microglial subtypes in mice with Aβ and tau pathologies mimic those seen in AD. *Tau4RΔK-AP* mice faithfully model not only the development of canonical AD pathologies, including the pathological conversion of wild-type mouse tau^24^, but also the disease stage-specific emergence of EADAM and LADAM in response to both Aβ plaques and tau deposition, emphasizing that our mouse model will prove useful in both analyzing AD disease mechanisms and testing therapeutic strategies for treating AD.

### Siglec Signaling in EADAM and LADAM

The identification of disease stage-specific EADAM and LADAM in our *Tau4RΔK-AP* model and AD brains raise the possibility that specific signaling pathways may govern the emergence of these microglial subtypes. CD33 (also known as Siglec-3)^48^, is one of the several microglial-specific genes that have been linked to AD susceptibility via genome-wide association studies (GWAS)^49^ and both mediates immune suppression by binding sialoglycan targets and attenuates phagocytosis by microglia during AD progression^48,50^ As a result, we first focused on characterizing the expression of sialic acid-binding immunoglobulin-like lectin (Siglec) family of proteins. Since human Siglecs belong to a family of 14 distinct transmembrane proteins, many of which are expressed in overlapping subsets of immune cells and inhibit immune activation^42,51^. In addition to Siglec-3, other members of the Siglec family could potentially regulate microglial function in AD. Our findings showing specific enrichment of Siglec-pathway genes in EADAM and LADAM strongly support this hypothesis. In particular, the observation that *Siglecg* is specifically expressed in LADAM while Siglec-10 (the human homolog of Siglec-G), is associated with late-stage AD pathologies exhibiting robust Aβ and Tau deposition, further supporting the view that Siglec signaling underlies the activation of LADAM in AD. Moreover, EADAM selectively expressed both *Siglec1* and *Siglecf*, while DAM and MHC Class II clusters both expressed *Siglecg*, indicating that Siglec-1, Siglec-F, and Siglec-G, respectively, represent potential biomarkers for detection of early- and late-stage disease. Taken together, our findings support the hypothesis that altered activity of disease stage-specific microglia during AD progression could be modulated by specific Siglec signaling pathways.

It is often difficult to identify Siglec orthologues between mice and humans, as this gene family evolved rapidly^51,52^. Mice do not express Siglec-8, and mouse Siglec-3 (mSiglec‑3) has different effector domains, binding specificity, cellular distribution and biological properties compared to those of human Siglec-3 (hSiglec‑3)^53,54^. In mouse microglia, immune inhibitory Siglec-F is among the most highly up-regulated genes (27-fold) in experimental neurodegenerative proteinopathy^55^. Siglec-F, a likely functional orthologue of Siglec-8^56^, is expressed in eosinophils as well as microglia^55,56^, binds to the same sialoglycans and sialoglycan mimetics as Siglec-8 (unlike mSiglec-3)^57,58^. Both Siglec-F and Siglec-8 also regulate eosinophilic inflammation^59,60^. Siglec-F was upregulated on a subset of reactive microglia in models of neurodegeneration, indicating the important role for Siglec-F/Siglec-8 in regulating microglial activation during neurodegeneration^61^. In this study, we observed that Siglec-F/8 expression emerged at earlier disease stages than Siglec-G/10 in both our mouse models and in AD. Our data also strongly suggest that Siglec-F is dependent on Aβ deposition, that Siglec-G is dependent on tau deposition, and that Aβ and tau deposition synergistically enhance the expression of both Siglecs. In addition, Siglec-G is enriched in microglia near myelinated regions, whereas Siglec-F+ microglia are more broadly expressed throughout the brain.

Our findings are consistent with a model whereby both Aβ and tau deposition are required to mediate the disease stage-specific induction of EADAM and LADAM through altered Siglec-dependent regulation of microglial function. This may offer new targets for the development of pre-symptomatic biomarkers and treatment for AD.

## Acknowledgments

This work was supported in part by the Maryland Stem Cell Research Fund (2019-MSCRFF-5124) to DWK, grants from the National Institute of Aging (R56AG068089) to RLS and TL, (R21AG073710) to TL, and the National Institute of Neurological Disorders and Stroke (R61NS115161) to PCW. We thank the Transcriptomics and Deep Sequencing Core (Johns Hopkins) for the sequencing of scRNA-Seq libraries and the Johns Hopkins ADRC Neuropathology Core for brain tissues.

## Conflicts of interest

The authors report no conflicts of interest.

---

**Figure S1.**
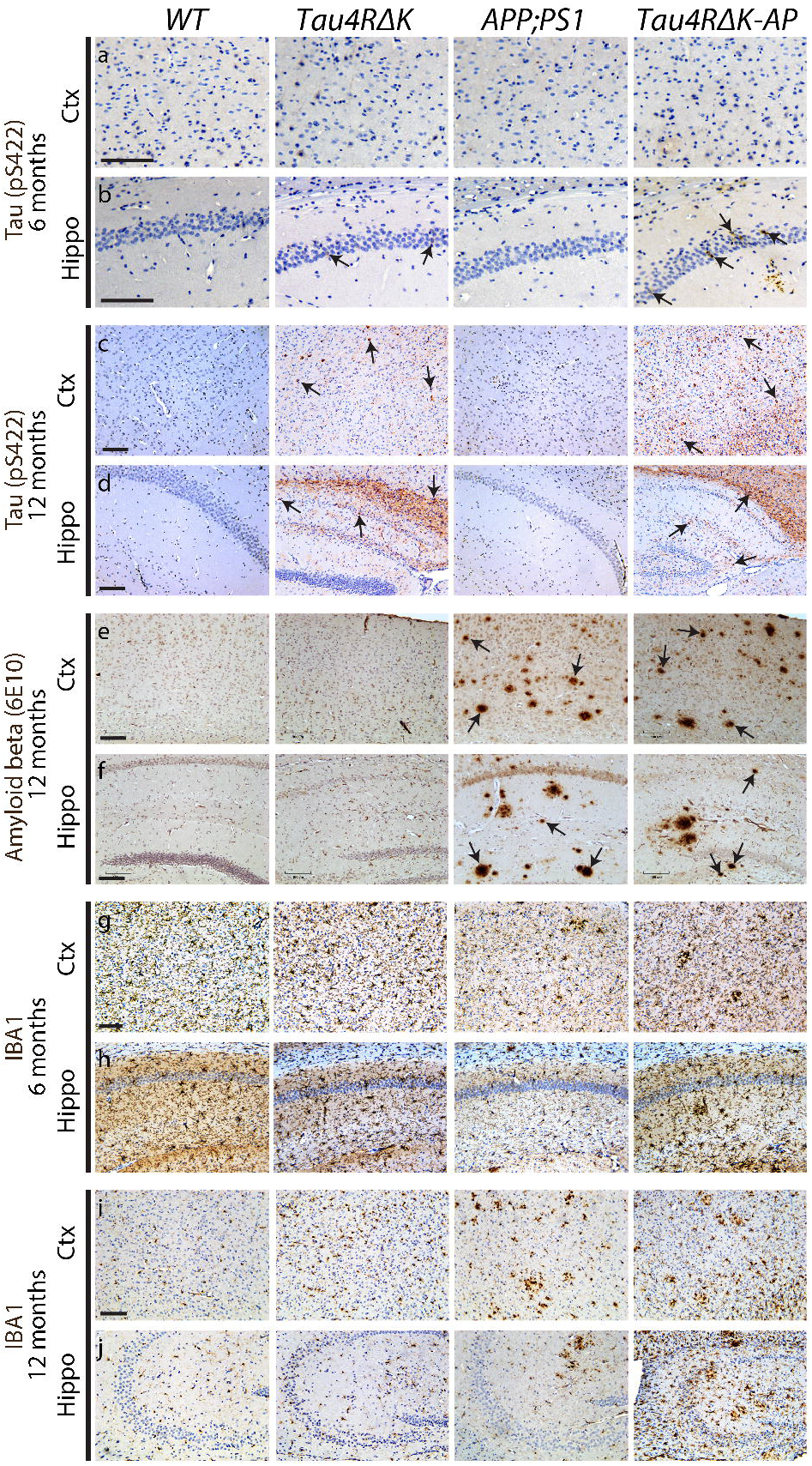
Histological validation of tau and Aβ pathologies in 6-month-old and 12-month-old *Tau4RΔK-AP* mice at. Immunostaining of tau (pS422) (a-d), and Aβ (6E10) (e, f), IBA1 (g-j) at 6-month-old (a, b, g, h) and 12-month-old (c-f, i, j); in the Ctx (a, c, e, g,i), and Hippo (b, d, f, h, j), in *WT*, *Tau4RΔK*, *APP;PS1*, *Tau4RΔK-AP* mice. Ctx = Cortex, Hippo = Hippocampus. Scale bars = 100 μm.

**Figure S2.**
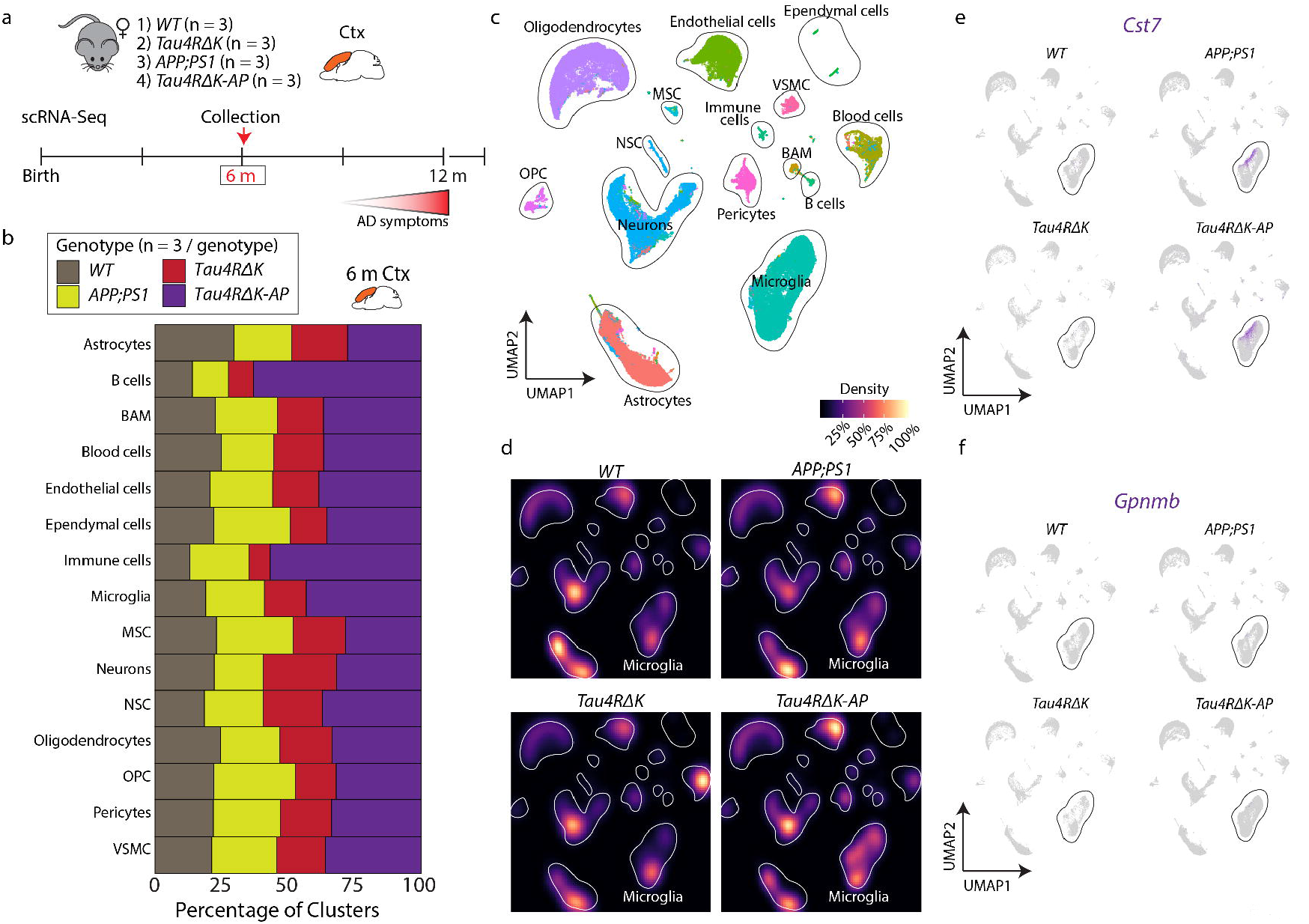
Distribution of cell types in 6-month-old cortex across genotypes. **a**, Schematic design of experiment: 4 genotypes (*WT, Tau4RΔK, APP:PS1, Tau4RΔK-AP*) in the cerebral cortex at 6-month-old and 12-month-old. **b**, Distribution of cell types across genotypes in the 6-month-old cortex. **c**, UMAP plot showing captured cell types in the 6-month-old cortex (all genotypes). **d**, UMAP plot showing the density of the captured cell types across genotypes in the 6-month-old cortex. **e**, UMAP plot showing DAM marker gene *Cst7*. **f**, UMAP plot showing DAM marker gene *Gpnmb*. BAM = brain-associated macrophages, Ctx = Cortex, MSC = muscle stem cells, OPC = oligodendrocyte precursor cells, VSMC = vascular smooth muscle cells.

**Figure S3.**
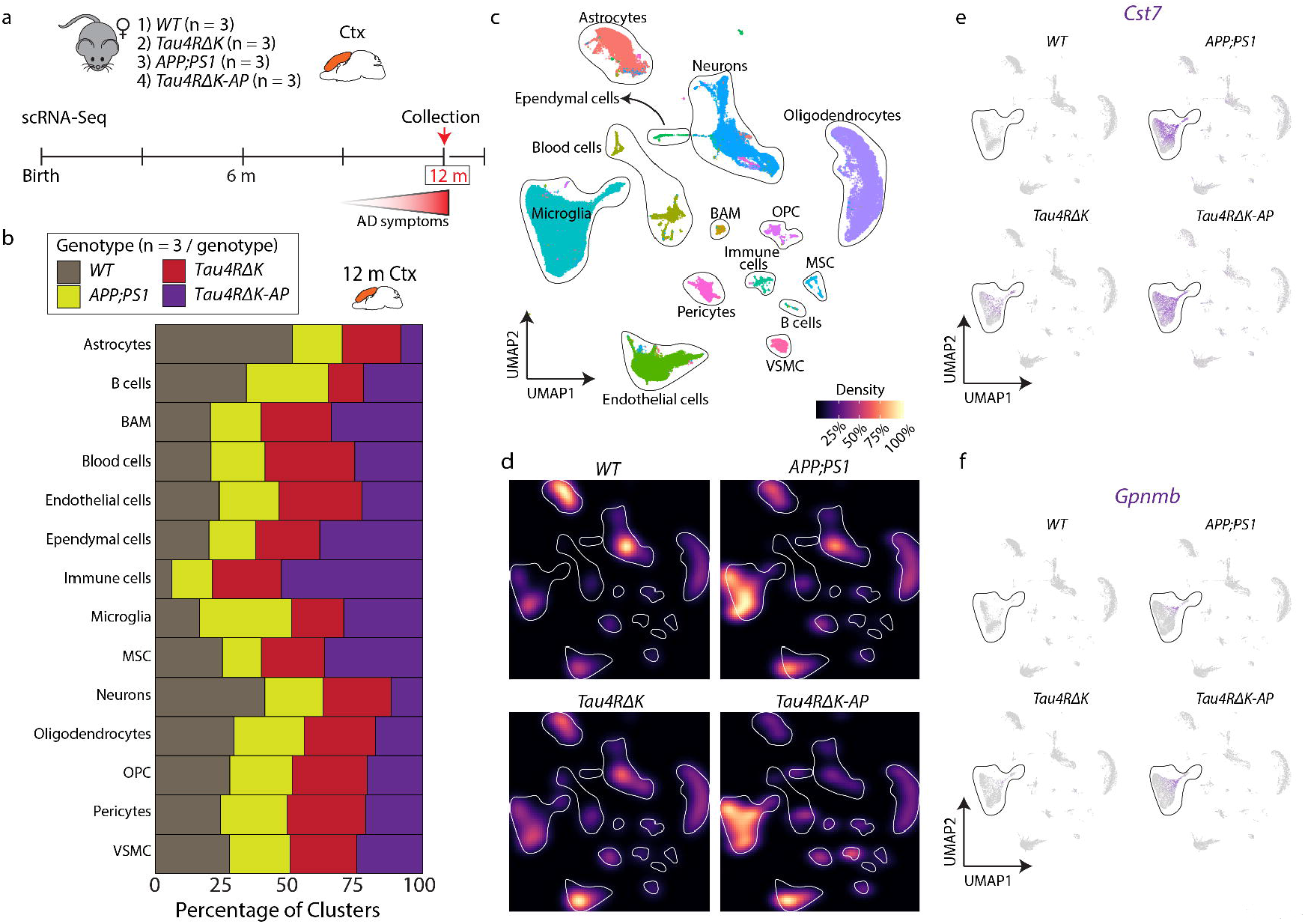
Distribution of cell types in 12-month-old cortex across genotypes. **a**, Schematic design of experiment: 4 genotypes (*WT, Tau4RΔK, APP:PS1, Tau4RΔK-AP*) in the cerebral cortex at 6 and 12-month-old. **b**, Distribution of cell types across genotypes in 12-month-old cortex. **c**, UMAP plot showing captured cell types in the 12-month-old cortex (all genotypes). **d**, UMAP plot showing the density of the captured cell types for each genotype in the 12-month-old cortex. **e**, UMAP plot showing DAM marker gene *Cst7*. **f**, UMAP plot showing DAM marker gene *Gpnmb*. BAM = brain-associated macrophages, Ctx = cortex, MSC = muscle stem cells, OPC = oligodendrocyte precursor cells, VSMC = vascular smooth muscle cells.

**Figure S4.**
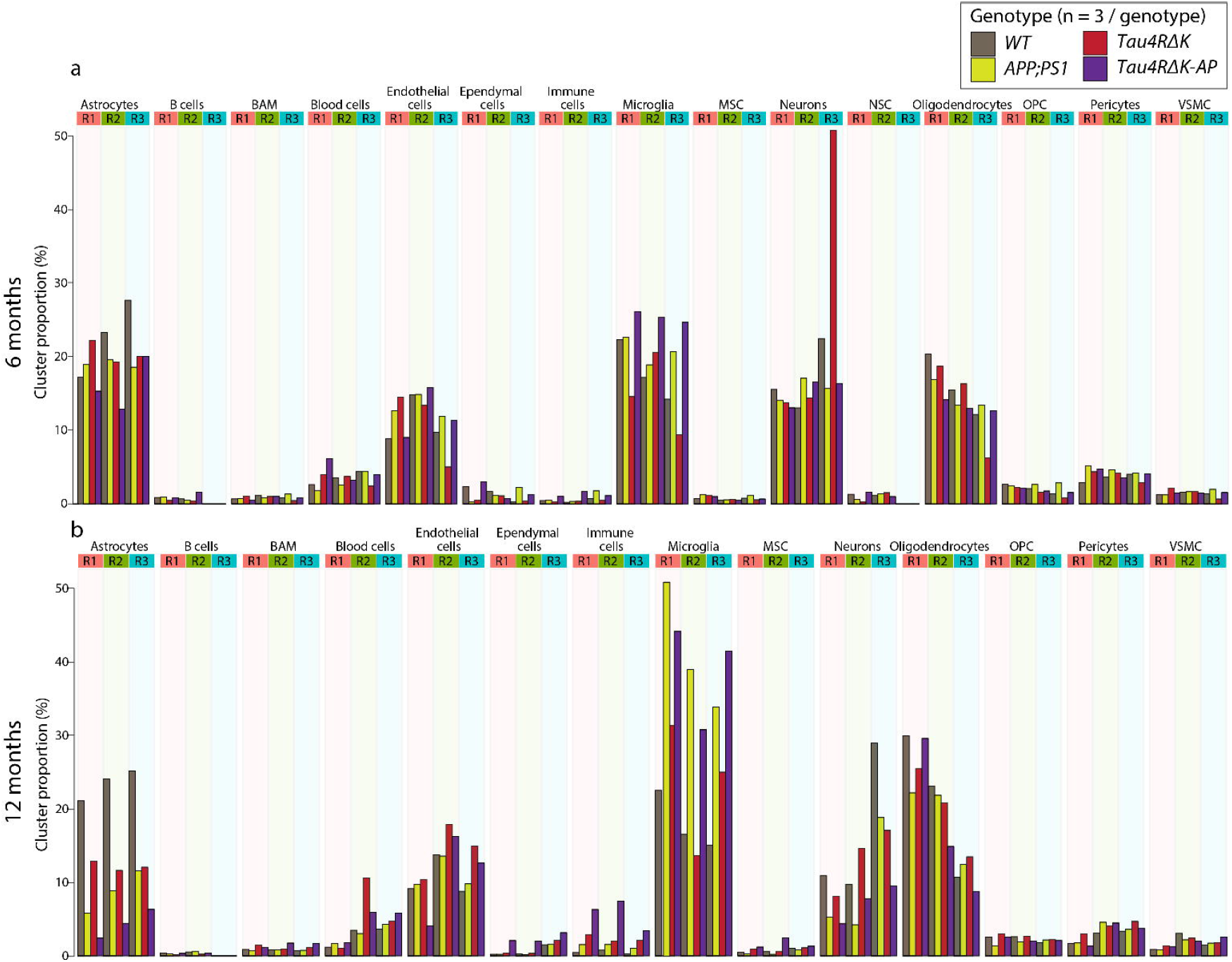
Distribution of clusters across genotypes. **a**, Distribution of cell types across 6-month-old scRNA-Seq triplicates. BAM = brain-associated macrophages, MSC = muscle stem cells, OPC = oligodendrocyte precursor cells, VSMC = vascular smooth muscle cells. **b**, Distribution of cell types across 12-month-old scRNA-Seq triplicates. BAM = brain-associated macrophages, MSC = muscle stem cells, OPC = oligodendrocyte precursor cells, VSMC = vascular smooth muscle cells.

**Figure S5.**
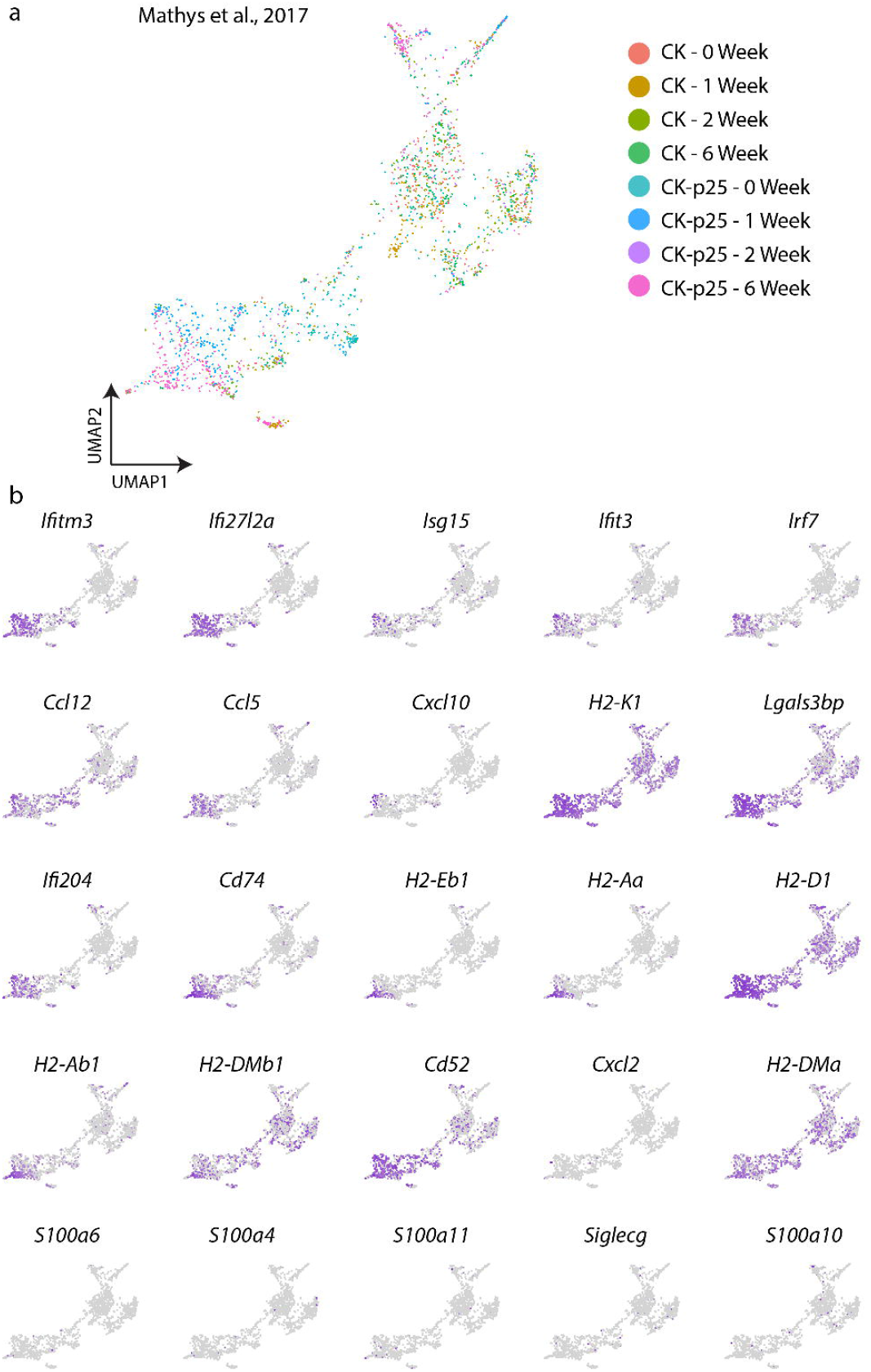
Analysis of EADAM and LADAM markers in other mouse models of neurodegenerative disease. **a**, UMAP plot showing CK-p25 neurodegenerative scNRA-Seq microglial dataset from Mathys et al., 2017. **b**, UMAP plot showing EADAM and LADAM-enriched genes. Note that while IFN and MHC genes are present but S100 genes and *Siglecg* are absent in the dataset.

**Figure S6.**
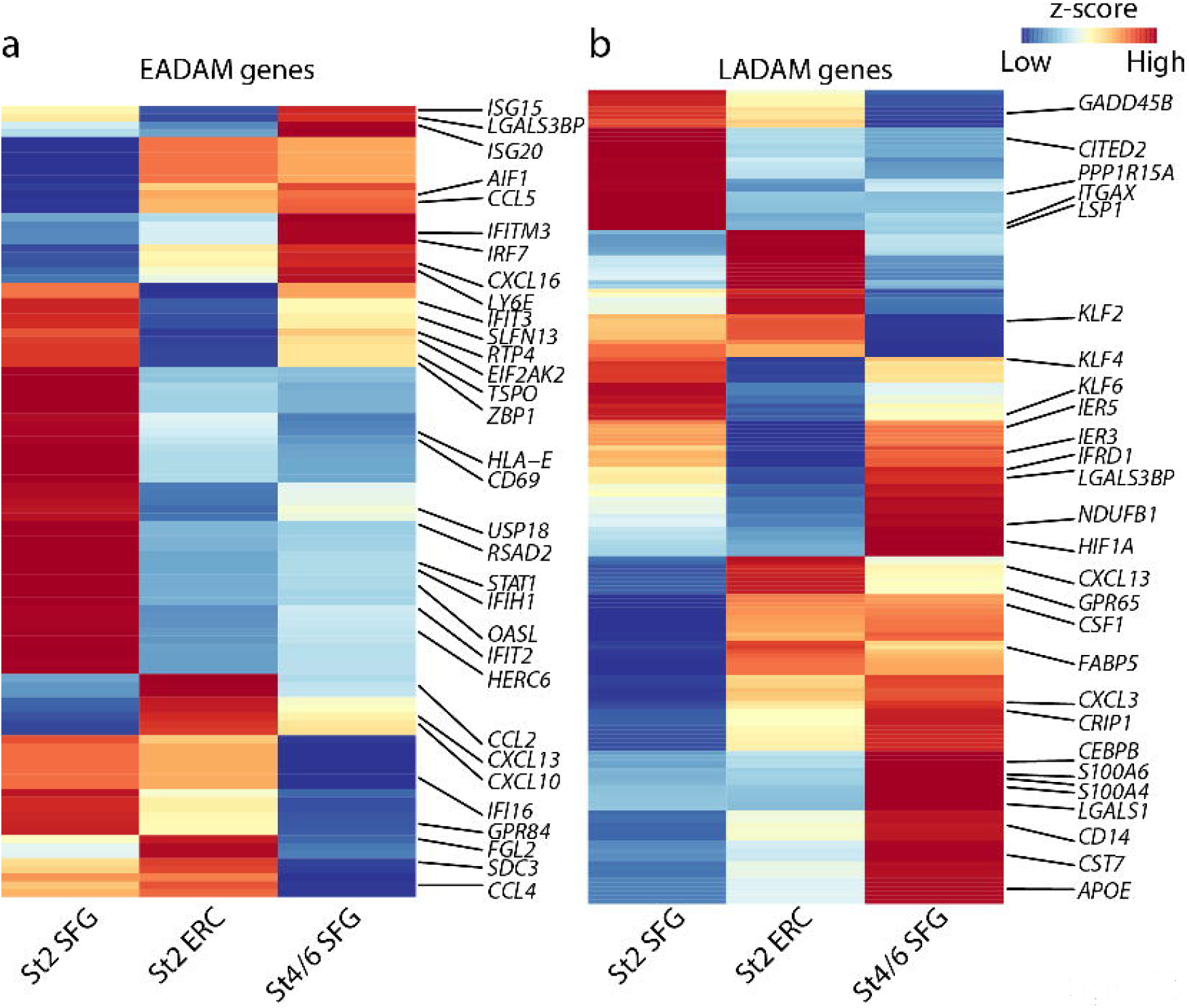
Microglial clusters in AD samples. Heatmap plot showing homologues of EADAM (a), and LADAM (b) genes in the human snRNA-Seq dataset.

**Figure S7.**
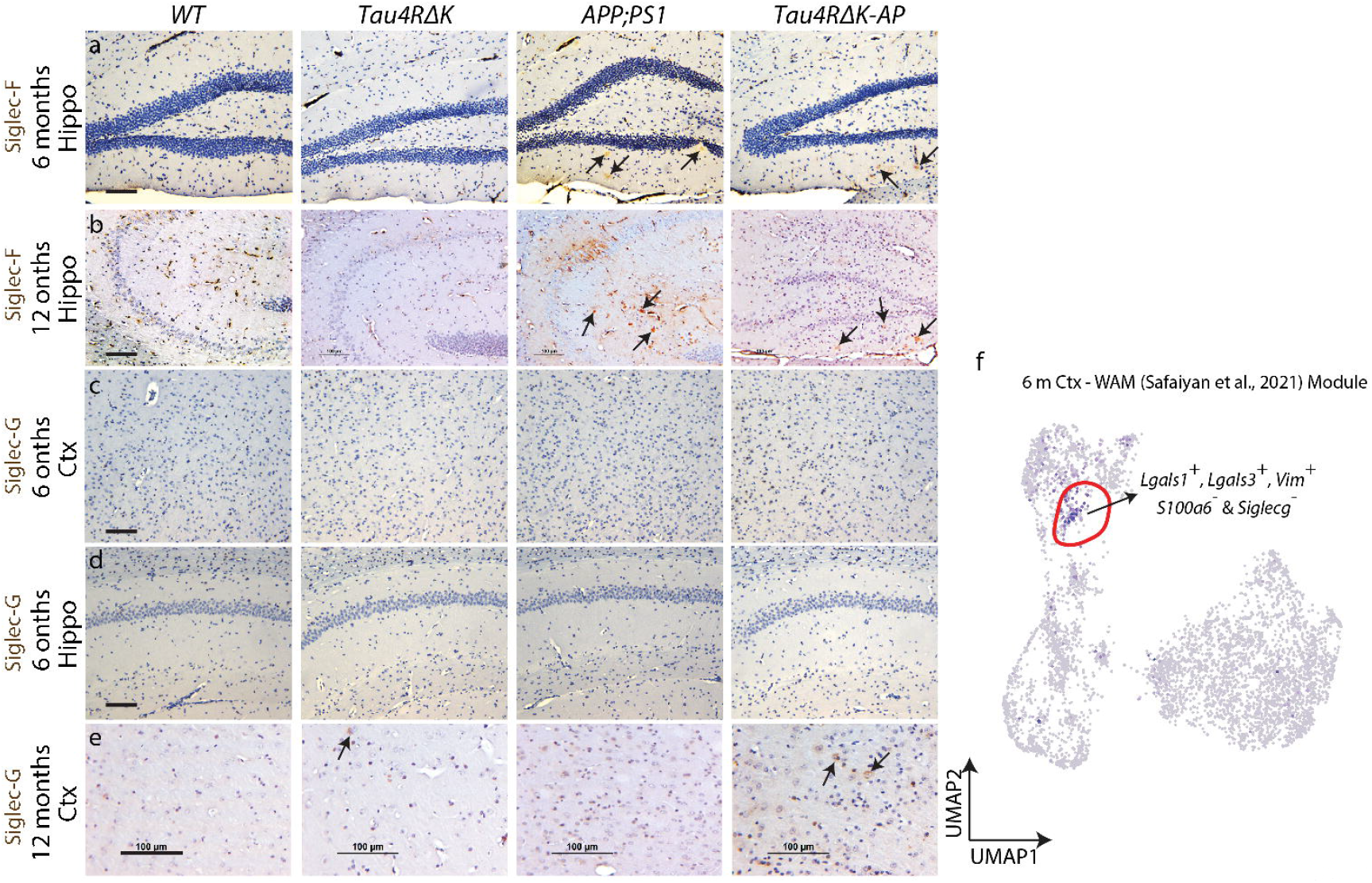
Histological validation of Siglec-F and Siglec-G across genotypes in 6 and 12-month-old mice. Immunostaining of Siglec-F (a, b), and Siglec-G (c-e), in *WT*, *Tau4RΔK*, *APP;PS1*, *Tau4RΔK-AP*; at 6-month-old (a, c) and 12-month-old (b, d, e); hippocampus (a, b, d), and the cortex (b, c, e). (f) UMAP plot of 6-month-old microglia, showing a small cluster (highlighted in red) that resembles molecular genes of WAM^25^. Scale bars = 100 μm.

**Figure S8.**
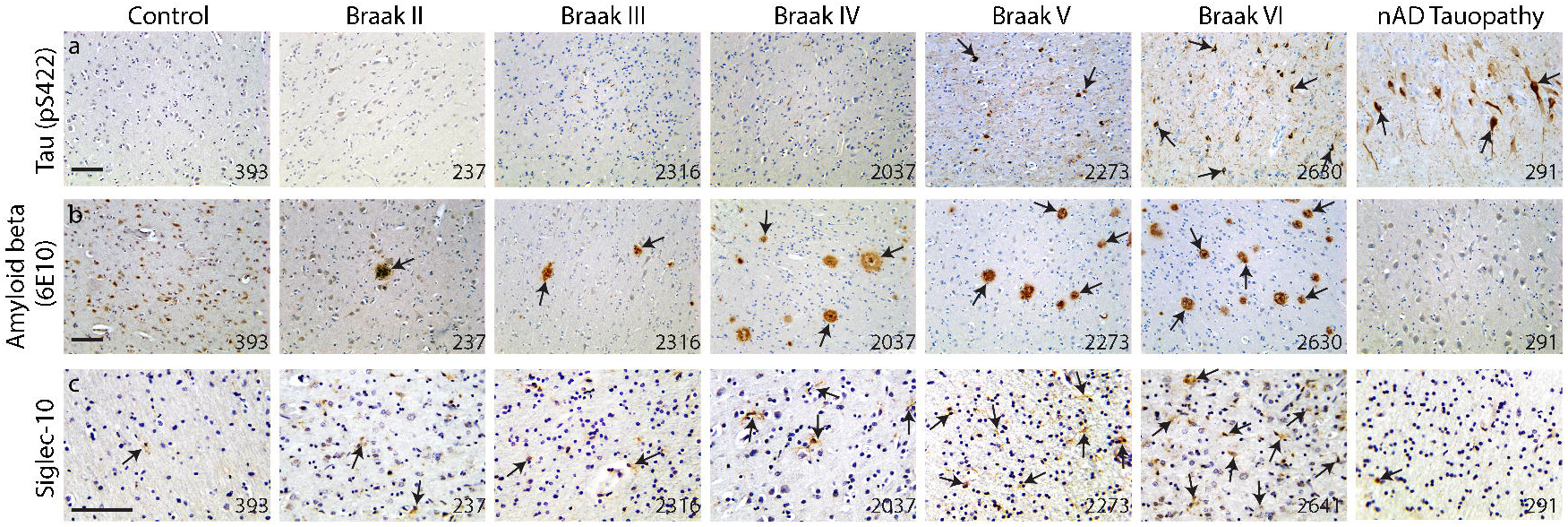
Histological validation of tau and Aβ pathologies and Siglec-10 during AD progression and in nAD Tauopathy. Immunostaining of Tau (pS422) (a), Aβ (6E10) (b), and Siglec-10 (c) in Control, Braak stage 2, Braak stage 3, Braak stage 4, Braak stage 5, Braak stage 6, and in nAD Tauopathy in the cortex. Numbers indicate BRC#. Scale bars = 100 μm.

## Materials and Methods

### Mice

*Tau4RΔK-AP* (*APP^swe^;PS1*D*E9;CamKII-tTA;TetO-TauRD*D*K*),*Tau4RΔK* (*CamKII-tTA;TetO-TauRDΔK*) and *APP^swe^;PS1ΔE9* transgenic mice were generated as described previously. *Tau4RDK* were generated by crossbreeding *TetO-TauRDΔK* transgenic mice carrying mutant *Tau* fragment with regulatory element *moPrP-tetP* promoter^63^ with *CamKII-tTA* mice to bring the Tau transgene under the control of the tet-off *CamKII* promoter^64^. *Tau4RΔK* mice were crossbred with *APP^swe^;PS1ΔE9* mice^26^ to generate *Tau4RΔK-AP* mice that develop both Tau pathology and Aβ amyloidosis. Because of sex differences observed in *Tau4RΔK-AP* mice, only female mice were used in this study. Brains were collected from the mice at 6-month-old and 12-month-old for scRNA-Seq, histological and immunohistological studies. To suppress the exogenous Tau expression during early development, mice were fed with Teklad Global 18% Protein Rodent Diet containing 200 mg/Kg doxycycline hydrochloride (Envigo Teklad Diets, Madison WI) during gestation and before weaning.

*CaMKIIα^CreERT2^;Tdp-43^F/F^* mice with *loxP* sites flanking *Tdp-43* exon 3 were generated as described previously^37^. Oral tamoxifen citrate was administered in the feed (Harlan Teklad) at an average 40 mg/kg/day for a 4-week period beginning at 6- or 9 month of age to induce recombination in excitatory forebrain neurons. Mice were singly housed during this period to monitor tamoxifen-feed intake. Afterwards, mice were returned to their original cage grouping with their littermates. Brain tissues were collected two month after the tamoxifen treatment for scRNA-Seq analysis.

All mice were housed in a climate-controlled facility (14-hour dark and 10-hour light cycle) with *ad libitum* access to food and water managed by Research Animal Resources (RAR) at Johns Hopkins University. All animal procedures were in accordance strictly with the National Institutes of Health Guide for the Care and Use of Laboratory Animals and were approved by the Johns Hopkins University Animal Care and Use Committee.

### scRNA-Seq Cell preparation

Mouse cortices were collected and dissociated using a previously published protocol^65,66^. Basically, cerebral cortices and hippocampi were dissected into Hibernate-A media with a 2% B-27 and GlutaMAX supplement (0.5 mM final). Tissues were dissociated in papain (Worthington) and debris was removed using OptiPrep density gradient media following cell dissociation. Cells were then processed immediately for scRNA-Seq.

### Study design and participant

The postmortem tissues used in the present study were provided by the Johns Hopkins Brain Resource Center. Formalin-fixed paraffin-embedded (FFPE) tissue sections (10 μm) of the inferior parietal region were obtained from 36 pathologically confirmed AD cases with Braak neurofibrillary stages II-VI and controls. All AD subjects had been prospectively recruited, clinically characterized by the Johns Hopkins Alzheimer’s Disease Research Center (ADRC), and underwent neuropathologic postmortem examination excluding Lewy body disease or non-AD tauopathies. The cohort (n= 36) included 22 males and 14 females ages 56 to 96 years (x= 77.58), 32 Whites, and 4 African-Americans. The average postmortem interval was 22 hours. In addition to the FFPE sections, we also examined frozen samples from the superior frontal gyrus and entorhinal cortex in two AD cases for snRNA-Seq. We also examined three FFPE hippocampal samples from patients with non-AD tauopathies. The clinical and autopsy components of this study were approved by the Johns Hopkins Medicine IRB.

### Isolation of nuclei from the frozen human brain for snRNA-Seq

Flash-frozen brain tissue was processed for snRNA-Seq following a modified 10x Genomics protocol. Briefly, lysis buffer containing (10 mM Tris-HCl pH7.4, 10 mM NaCl, 3 mM MgCl2, 0.1% Tween-20, 0.1% Nonidet P40, 0.01% Digitonin, 1 U/ul RNase inhibitor, 1% BSA) was added into a tube containing brain micropunch (~100 μm) and incubated on ice for 15 minutes with gentle pestle grinding (5 strokes every 3 min). Wash buffer (10 mM Tris-HCl pH7.4, 10 mM NaCl, 3 mM MgCl2, 0.1% Tween-20, 0.2 U/ul RNase inhibitor, 1% BSA) was added and filtered through a 50μm filter. Debris was removed using OptiPrep density gradient media and nuclei morphology was accessed under the light microscope. Nuclei were then immediately processed for snRNA-Seq.

### scRNA-Seq/snRNA-Seq generation

Cells or nuclei were loaded into the 10x Genomics Chromium Single Cell System (10x Genomics) and libraries were generated using v3 chemistry following the manufacturer’s instructions. Libraries were sequenced on Illumina NovaSeq6000. scRNA-Seq data were first processed through the Cell Ranger (v.3.1.0, 10x Genomics) with default parameters, aligned to the mm10 genome (refdata-cellranger-mm10-3.0.0), and matrix files were used for subsequent bioinformatic analysis. snRNA-Seq data were first processed through the Cell Ranger (v.5.0.0, 10x Genomics) with ‘include-introns’, aligned to the GRCh38 genome (refdata-gex-GRCh38-2020-A), and matrix files were used for subsequent bioinformatic analysis

### scRNA-Seq data analysis

#### 1. Processing scRNA-Seq datasets

Seurat v3.15^67^ was used to process matrix files, first by selecting cells with more than 500 genes, 1000 UMI, and less than 50% mitochondrial genes and 20% ribosomal genes. Datasets were then normalized using Seurat ‘scTransform’ function with regressing the number of genes and UMIs using ‘vars.to.regress’, and Harmony v1.0^68^ was used to adjust for batch variation by treating individual scRNA-Seq run as a variance group.

The top 20 variables obtained from Harmony analysis were used for UMAP dimensional reduction. The Louvain clustering algorithm was used to first identify main cell types with default resolution and the top 20 reduction variables obtained from Harmony analysis. Individual cell types were first identified by cross-referencing to previous scRNA-Seq datasets^66,69^ and ASCOT. Doublet cells - cells that express both cell-type-specific genes (~5% of the overall datasets) and unhealthy cells (when mitochondrial/ribosomal genes were expressed at much higher levels compared to other clusters) were excluded from further analysis. This was done in individual ages, in all genotypes. Individual microglia were subsetted and processed as described above. Any doublet cells (~2%) that couldn’t be previously identified were removed for further analysis. The combination of 6-month-old and 12-month-old microglia, Homeostatic 2 (microglia cluster that expresses high IEGs due to technical artifacts) were removed. Enriched microglia cluster genes in our datasets, pseudo-bulked enriched genes in previously published human datasets^40^, or WAM LADAM genes were superimposed using Seurat *AddModuleScore* function and module score was then calculated.

Differential gene tests on the scRNA-Seq datasets were initially performed using Seurat v3.15 *FindAllMarkers* using ‘LR’ with default parameters on all expressed genes and using gene numbers, UMIs, and cell numbers as variables (‘latent.vars’).

Two initial microglia clusters expressing high levels of homeostatic microglial genes, such as *Tmem119* and *P2ry12*, were distributed almost equally among all four genotypes (Fig.2a-c), but one cluster showed higher expression of immediate-early genes such as *Fos, Atf3*, and *Junb*. This was detected across genotypes and derived from technical artifacts during the heat-activated enzymatic dissociation step^70^ and these microglia were then removed for any downstream analysis.

#### 2. Regulons

To identify regulons controlling gene expression in different microglial populations across ages and genotypes, SCENIC^62^ using python implemented pySCENIC v.0.10.0 Gene regulatory networks (using --masks_dropouts), regulons and network activity of regulons were calculated using default parameters with mm10 feather files on the microglial dataset using raw count matrix. Regulon specificity scores were ranked following the SCENIC pipeline and top regulons with z-score higher than 2 were identified as microglia-specific.

#### 3. GO pathway analysis

Top differential genes (adjusted p-value < 0.05, fold change > 0.5) in each microglial cluster were used as an input for GO pathway analysis using ClusterProfiler (v3.12.0)^71^.

#### 4. RNA velocity

RNA velocity^72^ was utilized to understand the dynamic state of microglia, and how each genotype-specific disease-associated microglia cluster was initiated across AD progression. Kallisto v0.46.2 and bustools v2.27.9^73,74^ python wrapper kb-python was used to obtain spliced and unspliced transcripts using --lamanno with GRCm38 mouse genome. Scanpy v1.5.1^75^ and scVelo v0.2.1^76^ were used to process the Kallisto output with default parameters, based on UMAP coordinates obtained from Seurat.

#### 5. Pseudotime analysis

Monocle v3.0.2^77^ was used to perform pseudotime analysis to identify differences in gene expression across microglial disease states, where the trajectory routes used for pseudotime analysis were identified based on trajectories from RNA velocity analysis, and high-variance genes (q < 0.001) were used for pseudotime plotting.

### Histology and Immunohistochemical Analysis

For the histological and immunohistochemical analysis, mice were anesthetized and brains were removed and weighed. Hemibrains were fixed by submerging into 4% PFA in PBS, embedded into the paraffin, and sectioned in the sagittal plane. For histological analyses, 10 μm brain sections were stained with hematoxylin and eosin (H&E) or Cresyl violet. For immunohistochemical analysis, sections were treated with 10 mM citrate buffer (pH 6.0) by microwave in high power for 6 minutes for antigen retrieval; endogenous peroxidase was quenched by treating with 0.3% H_2_O_2_, and nonspecific binding of antibodies was eliminated using blocking buffer (10% normal goat serum in PBS with 0.3% Triton-X) for one hour at room temperature. The primary antibody was prepared in a blocking buffer and was applied overnight at 4°C, followed by a secondary antibody for 30 min incubation at room temperature. For the secondary antibody and avidin-biotinylated peroxidase system, we used the Vectastain Universal Elite ABC kit (Vector Laboratories). Brain sections were stained with: antiserum against Aβ peptides 6E10 (1:1,000; SIG-39300, Covance); rabbit antiserum against phosphorylated S422 of tau (1:2,000; 44764G, Invitrogen, Carlsbad, CA); polyclonal antiserum against GFAP (1:1,000, Z0334, Dako Corporation, Carpinteria, CA); polyclonal antiserum against microglial (IBA1, 1:1,000, CP290, Biocare Medical, CA); polyclonal antiserum against Siglec-10 (1:50, HPA027093, Sigma); Monoclonal Antibody Siglec-F (CD170, 1:300, 14-1702-82, ThermoFisher Scientific). Quantitative score of Siglec10 positive cells in the human brains was the average counts of two brain sections in a power (2,000X) microscope field using the ImageJ program. All data were analyzed statistically by one-way ANOVA with post hoc t-test for multiple comparisons using Excel. In all tests, values of *p < 0.05* were considered to indicate significance.

## Notes

### Competing Interest Statement

The authors have declared no competing interest.

### Summary of Updates

Figure 5 was not uploaded previously.

## References

1. Bohlen, C. J., Friedman, B. A., Dejanovic, B. & Sheng, M. Microglia in Brain Development, Homeostasis, and Neurodegeneration. Annu. Rev. Genet. 53, 263–288 (2019).

2. Hickman, S., Izzy, S., Sen, P., Morsett, L. & El Khoury, J. Microglia in neurodegeneration. Nat. Neurosci. 21, 1359–1369 (2018).

3. 2021 Alzheimer’s disease facts and figures. Alzheimers. Dement. 17, 327–406 (2021).

4. Bekris, L. M., Yu, C.-E., Bird, T. D. & Tsuang, D. W. Genetics of Alzheimer disease. J. Geriatr. Psychiatry Neurol. 23, 213–227 (2010).

5. Saunders, A. M. et al. *Association of apolipoprotein E allele epsilon 4 with late-onset* familial and sporadic Alzheimer’s disease. Neurology 43, 1467–1472 (1993).

6. Farrer, L. A. et al. Effects of age, sex, and ethnicity on the association between apolipoprotein E genotype and Alzheimer disease. A meta-analysis. APOE and Alzheimer Disease Meta Analysis Consortium. JAMA 278, 1349–1356 (1997).

7. Guerreiro, R. et al. TREM2 variants in Alzheimer’s disease. N. Engl. J. Med. 368, 117–127 (2013).

8. Jonsson, T. et al. Variant of TREM2 associated with the risk of Alzheimer’s disease. N. Engl. J. Med. 368, 107–116 (2013).

9. Sims, R. et al. *Rare coding variants in PLCG2, ABI3, and TREM2 implicate microglial-* mediated innate immunity in Alzheimer’s disease. Nat. Genet. 49, 1373–1384 (2017).

10. Hollingworth, P. et al. Common variants at ABCA7, MS4A6A/MS4A4E, EPHA1, CD33 and CD2AP are associated with Alzheimer’s disease. Nat. Genet. 43, 429–435 (2011).

11. Naj, A. C. et al. Common variants at MS4A4/MS4A6E, CD2AP, CD33 and EPHA1 are associated with late-onset Alzheimer’s disease. Nat. Genet. 43, 436–441 (2011).

12. Butovsky, O. & Weiner, H. L. Microglial signatures and their role in health and disease. Nat. Rev. Neurosci. 19, 622–635 (2018).

13. Heneka, M. T. et al. Neuroinflammation in Alzheimer’s disease. Lancet Neurol. 14, 388–405 (2015).

14. Hamelin, L. et al. *Early and protective microglial activation in Alzheimer’s disease: a* prospective study using 18F-DPA-714 PET imaging. Brain 139, 1252–1264 (2016).

15. Ising, C. et al. NLRP3 inflammasome activation drives tau pathology. Nature 575, 669–673 (2019).

16. Fani Maleki, A. & Rivest, S. Innate Immune Cells: Monocytes, Monocyte-Derived Macrophages and Microglia as Therapeutic Targets for Alzheimer’s Disease and Multiple Sclerosis. Front. Cell. Neurosci. 13, 355 (2019).

17. Masuda, T., Sankowski, R., Staszewski, O. & Prinz, M. Microglia Heterogeneity in the Single-Cell Era. Cell Rep. 30, 1271–1281 (2020).

18. Xue, F. & Du, H. TREM2 Mediates Microglial Anti-Inflammatory Activations in Alzheimer’s Disease: Lessons Learned from Transcriptomics. Cells 10, (2021).

19. Wes, P. D., Holtman, I. R., Boddeke, E. W. G. M., Möller, T. & Eggen, B. J. L. Next generation transcriptomics and genomics elucidate biological complexity of microglia in health and disease. Glia 64, 197–213 (2016).

20. Keren-Shaul, H. et al. A Unique Microglia Type Associated with Restricting Development of Alzheimer’s Disease. Cell 169, 1276–1290.e17 (2017).

21. Li, Q. et al. Developmental Heterogeneity of Microglia and Brain Myeloid Cells Revealed by Deep Single-Cell RNA Sequencing. Neuron 101, 207–223.e10 (2019).

22. Masuda, T. et al. *Spatial and temporal heterogeneity of mouse and human microglia at* single-cell resolution. Nature 566, 388–392 (2019).

23. Zhou, Y. et al. *Human and mouse single-nucleus transcriptomics reveal TREM2-* dependent and TREM2-independent cellular responses in Alzheimer’s disease. Nat. Med. 26, 131–142 (2020).

24. Li, T. et al. *The neuritic plaque facilitates pathological conversion of tau in an* Alzheimer’s disease mouse model. Nature Communications vol. 7 (2016).

25. Safaiyan, S. et al. White matter aging drives microglial diversity. Neuron 109, 1100–1117.e10 (2021).

26. Jankowsky, J. L., Xu, G., Fromholt, D., Gonzales, V. & Borchelt, D. R. Environmental enrichment exacerbates amyloid plaque formation in a transgenic mouse model of Alzheimer disease. J. Neuropathol. Exp. Neurol. 62, 1220–1227 (2003).

27. Sierksma, A. et al. Novel Alzheimer risk genes determine the microglia response to amyloid-β but not to TAU pathology. doi:10.1101/491902.

28. Frigerio, C. S. et al. *The Major Risk Factors for Alzheimer’s Disease: Age, Sex, and* Genes Modulate the Microglia Response to Aβ Plaques. Cell Reports vol. 27 1293–1306.e6 (2019).

29. Yang, H. S. et al. Natural genetic variation determines microglia heterogeneity in wild-derived mouse models of Alzheimer’s disease. doi:10.1101/2020.06.02.130237.

30. Roy, E. R. et al. *Type I interferon response drives neuroinflammation and synapse loss* in Alzheimer disease. J. Clin. Invest. 130, 1912–1930 (2020).

31. Bryan, K. J. et al. Expression of CD74 is increased in neurofibrillary tangles in Alzheimer’s disease. Molecular Neurodegeneration vol. 3 13 (2008).

32. Mathys, H. et al. Temporal Tracking of Microglia Activation in Neurodegeneration at Single-Cell Resolution. Cell Rep. 21, 366–380 (2017).

33. Meilandt, W. J. et al. Trem2 Deletion Reduces Late-Stage Amyloid Plaque Accumulation, Elevates the Aβ42:Aβ40 Ratio, and Exacerbates Axonal Dystrophy and Dendritic Spine Loss in the PS2APP Alzheimer’s Mouse Model. J. Neurosci. 40, 1956–1974 (2020).

34. Lee, S.-H. et al. *Trem2 restrains the enhancement of tau accumulation and* neurodegeneration by β-amyloid pathology. Neuron 109, 1283–1301.e6 (2021).

35. Götz, J., Chen, F., Barmettler, R. & Nitsch, R. M. Tau filament formation in transgenic mice expressing P301L tau. J. Biol. Chem. 276, 529–534 (2001).

36. Ozmen, L., Albientz, A., Czech, C. & Jacobsen, H. Expression of transgenic APP mRNA is the key determinant for beta-amyloid deposition in PS2APP transgenic mice. Neurodegener. Dis. 6, 29–36 (2009).

37. LaClair, K. D. et al. Depletion of TDP-43 decreases fibril and plaque β-amyloid and exacerbates neurodegeneration in an Alzheimer’s mouse model. Acta Neuropathol. 132, 859–873 (2016).

38. Morabito, S. et al. Single-nucleus chromatin accessibility and transcriptomic characterization of Alzheimer’s disease. Nat. Genet. 53, 1143–1155 (2021).

39. Pande, R. et al. Single Cell Atlas of Human Putamen Reveals Disease Specific Changes in Synucleinopathies: Parkinson’s Disease and Multiple System Atrophy. doi:10.1101/2021.05.06.442950.

40. Leng, K. et al. Molecular characterization of selectively vulnerable neurons in Alzheimer’s Disease. doi:10.1101/2020.04.04.025825.

41. Siddiqui, S. S. et al. Siglecs in Brain Function and Neurological Disorders. Cells 8, (2019).

42. Macauley, M. S., Crocker, P. R. & Paulson, J. C. Siglec-mediated regulation of immune cell function in disease. Nat. Rev. Immunol. 14, 653–666 (2014).

43. Leng, F. & Edison, P. Neuroinflammation and microglial activation in Alzheimer disease: where do we go from here? Nat. Rev. Neurol. 17, 157–172 (2021).

44. Grubman, A. et al. *Transcriptional signature in microglia associated with Aβ plaque* phagocytosis. Nat. Commun. 12, 3015 (2021).

45. Leyns, C. E. G. et al. TREM2 function impedes tau seeding in neuritic plaques. Nat. Neurosci. 22, 1217–1222 (2019).

46. Mathys, H. et al. Single-cell transcriptomic analysis of Alzheimer’s disease. Nature 570, 332–337 (2019).

47. Marschallinger, J. et al. *Lipid-droplet-accumulating microglia represent a dysfunctional* and proinflammatory state in the aging brain. Nature Neuroscience vol. 23 194–208 (2020).

48. Bradshaw, E. M. et al. *CD33 Alzheimer’s disease locus: altered monocyte function and* amyloid biology. Nat. Neurosci. 16, 848–850 (2013).

49. Podleśny-Drabiniok, A., Marcora, E. & Goate, A. M. Microglial Phagocytosis: A Disease-Associated Process Emerging from Alzheimer’s Disease Genetics. Trends Neurosci. 43, 965–979 (2020).

50. Griciuc, A. et al. *Alzheimer’s disease risk gene CD33 inhibits microglial uptake of* amyloid beta. Neuron 78, 631–643 (2013).

51. Varki, A., Schnaar, R. L. & Crocker, P. R. I-Type Lectins. in *Essentials of Glycobiology* (eds. Varki, A. et al.) (Cold Spring Harbor Laboratory Press, 2017).

52. Varki, A. Multiple changes in sialic acid biology during human evolution. Glycoconj. J. 26, 231–245 (2009).

53. Bhattacherjee, A. et al. Repression of phagocytosis by human CD33 is not conserved with mouse CD33. Commun Biol 2, 450 (2019).

54. Brinkman-Van der Linden, E. C. M. et al. *CD33/Siglec-3 binding specificity, expression* pattern, and consequences of gene deletion in mice. Mol. Cell. Biol. 23, 4199–4206 (2003).

55. Lunnon, K. et al. *Systemic inflammation modulates Fc receptor expression on microglia* during chronic neurodegeneration. J. Immunol. 186, 7215–7224 (2011).

56. Bochner, B. S. Siglec-8 on human eosinophils and mast cells, and Siglec-F on murine eosinophils, are functionally related inhibitory receptors. Clin. Exp. Allergy 39, 317–324 (2009).

57. Yu, H. et al. *Siglec-8 and Siglec-9 binding specificities and endogenous airway ligand* distributions and properties. Glycobiology 27, 657–668 (2017).

58. Nycholat, C. M. et al. *A Sulfonamide Sialoside Analogue for Targeting Siglec-8 and -F* on Immune Cells. J. Am. Chem. Soc. 141, 14032–14037 (2019).

59. McMillan, S. J., Richards, H. E. & Crocker, P. R. Siglec-F-dependent negative regulation of allergen-induced eosinophilia depends critically on the experimental model. Immunol. Lett. 160, 11–16 (2014).

60. Zhang, M. et al. *Defining the in vivo function of Siglec-F, a CD33-related Siglec* expressed on mouse eosinophils. Blood 109, 4280–4287 (2007).

61. Morshed, N. et al. Phosphoproteomics identifies microglial Siglec-F inflammatory response during neurodegeneration. Mol. Syst. Biol. 16, e9819 (2020).

62. Aibar, S. et al. SCENIC: single-cell regulatory network inference and clustering. Nat. Methods 14, 1083–1086 (2017).

63. Jankowsky, J. L. et al. *Persistent amyloidosis following suppression of Abeta production* in a transgenic model of Alzheimer disease. PLoS Med. 2, e355 (2005).

64. Mayford, M. et al. Control of memory formation through regulated expression of a CaMKII transgene. Science 274, 1678–1683 (1996).

65. Kim, D. W. et al. Gene regulatory networks controlling differentiation, survival, and diversification of hypothalamic Lhx6-expressing GABAergic neurons. Commun Biol 4, 95 (2021).

66. Kim, D. W. et al. The cellular and molecular landscape of hypothalamic patterning and differentiation from embryonic to late postnatal development. Nat. Commun. 11, 4360 (2020).

67. Stuart, T. et al. Comprehensive Integration of Single-Cell Data. Cell vol. 177 1888–1902.e21 (2019).

68. Korsunsky, I. et al. Fast, sensitive and accurate integration of single-cell data with Harmony. Nat. Methods 16, 1289–1296 (2019).

69. Yoo, S., Cha, D., Kim, D. W., Hoang, T. V. & Blackshaw, S. Tanycyte-Independent Control of Hypothalamic Leptin Signaling. Front. Neurosci. 13, 240 (2019).

70. Marsh, S. E. et al. Single Cell Sequencing Reveals Glial Specific Responses to Tissue Processing & Enzymatic Dissociation in Mice and Humans. doi:10.1101/2020.12.03.408542.

71. Yu, G., Wang, L.-G., Han, Y. & He, Q.-Y. clusterProfiler: an R Package for Comparing Biological Themes Among Gene Clusters. OMICS: A Journal of Integrative Biology vol. 16 284–287 (2012).

72. La Manno, G. et al. RNA velocity of single cells. Nature 560, 494–498 (2018).

73. Melsted, P. et al. Modular and efficient pre-processing of single-cell RNA-seq. doi:10.1101/673285.

74. Bray, N. L., Pimentel, H., Melsted, P. & Pachter, L. Near-optimal probabilistic RNA-seq quantification. Nat. Biotechnol. 34, 525–527 (2016).

75. Alexander Wolf, F., Angerer, P. & Theis, F. J. SCANPY : large-scale single-cell gene expression data analysis. Genome Biol. 19, 15 (2018).

76. Bergen, V., Lange, M., Peidli, S., Alexander Wolf, F. & Theis, F. J. *Generalizing RNA velocity to transient cell states through dynamical modeling*. doi:10.1101/820936.

77. Qiu, X. et al. Single-cell mRNA quantification and differential analysis with Census. Nat. Methods 14, 309–315 (2017).

